# Microbiome and Metabolome driven differentiation of TGF-β producing Tregs leads to Senescence and HIV latency

**DOI:** 10.1101/2020.12.15.422949

**Authors:** Khader Ghneim, Ashish Arunkumar Sharma, Susan Pereira Ribeiro, Slim Fourati, Jeffrey Ahlers, Deanna Kulpa, Xuan Xu, Jessica Brehm, Aarthi Talla, Sahaana Arumugam, Samuel Darko, Benigno Rodriguez, Carey Shive, Razvan Cristescu, Andrey Loboda, Robert Balderas, I-Ming Wang, Peter Hunt, Daniel Lamarre, Daniel Douek, Daria Hazuda, Michael M. Lederman, Steven Deeks, Rafick-Pierre Sekaly

**Author notes:** Both authors equally contributed to the manuscript (Khader Ghneim, Ashish Arunkumar Sharma). Rafick-Pierre Sekaly and Ashish Arunkumar Sharma are the corresponding authors for this manuscript. Please direct all inquiries to. Deceased (Benigno Rodriguez).

## Abstract

Current therapeutic interventions to eradicate latent HIV (“reservoir”) and restore immune function in ART-treated HIV infection have yet to show efficacy. To explore mechanisms of HIV persistence, we apply an integrated systems biology approach and identify a distinct group of individuals with poor CD4 T-cell reconstitution (Immunologic non-responders, “INRs”) and high frequencies of cells with inducible HIV. Contrary to the prevailing notion that immune activation drives HIV persistence and immune dysfunction, peripheral blood leukocytes from these subjects have enhanced expression of a network of genes regulated by cellular senescence driving transcription factors (TFs) FOXO3, SMAD2 and IRF3. In these subjects, increased frequencies of regulatory T cells and expression of the TGF-β signaling cascade are complimented by the downregulation of cell cycle, metabolic and pro-inflammatory pathways. Lactobacillaceae family and metabolites (members of the butyrate family – i.e. α-ketobutyrate) were correlated with Treg frequencies in “Senescent-INRs” *ex vivo,* triggered the differentiation of TGF-β producing Tregs and promoted HIV latency establishment *in vitro.* These cascades, downstream of PD-1/TGF-β, prevent memory T cell differentiation and are associated with an increase in frequencies of cells with inducible HIV *ex vivo.* Our findings identify cellular senescence responses that can be targeted by PD-1 or TGF-β specific interventions that have shown safety and efficacy in cancer, and may prove to be crucial for HIV eradication.

## Introduction

HIV infection remains a major global public health concern (WHO, https://www.who.int/news-room/fact-sheets/detail/hiv-aids). Although effective combined anti-retroviral therapy (cART) has altered the course of HIV disease by preventing viral replication, it is not curative and does not fully restore immune function^1^. Depending on the population studied and outcome definitions^2^, up to 20% of cART-treated subjects fail to reconstitute CD4+ T-cell numbers despite years of effective treatment with sustained HIV suppression, and are predisposed to excessive risk of non-HIV comorbidities^3^.

These immune non-responder subjects (INRs) show severe homeostatic alterations in CD4 T-cells, including lower frequencies of naïve CD4 T-cells, accumulation of highly differentiated T-cells, T-cell activation and apoptosis^4–6^. Impairment in IL-7/L-7R signal transduction axis^7^, heightened chronic inflammation and persistent type I interferon production; are thought to be important mediators of poor CD4 T cell homeostasis and reduced thymic function^8^ in the INRs. Additionally, CD4 T cells in the INRs have higher expression of replicative senescence^9^ and exhaustion markers^10^, and show an increase in frequencies of immunosuppressive CD4+ regulatory T-cells (Tregs)^11^. Using multivariate analyses, we previously reported that naive T-cell depletion, higher frequencies of cycling CD4+ central memory (CM) and effector memory (EM) T-cells, expression of the T-cell activation markers (CD38, HLA-DR, CCR5 and/or PD-1 (p<0.0001)), and levels of soluble CD14 (sCD14) distinguished INRs from immune responder subjects (IRs). These groups are distinguishable even when adjusted for CD4+ T-cell nadir, age at cART initiation, and other clinical indices^12–14^. Collectively these studies highlight the contribution of dysregulated pro-inflammatory pathways to the lack of immune reconstitution in INRs.

Despite the ability of cART to inhibit HIV replication, the proviral DNA integrates into the human genome and persists within memory CD4 T-cells^15^. This cellular “reservoir” reignites rounds of virus replication if cART is interrupted^16^, stressing the need for long-term cART administration. Indeed, reconstitution of CD4 counts during uninterrupted cART leads to a decrease in intact pro-viral DNA^17^, whereas elevated levels of inflammatory markers characteristic of INRs (IL6, TNF□, IL1β, IFN□/β, MIP1□/β, RANTES) are associated with HIV persistence^18^. The activated state of the innate immune system can trigger HIV replication, alter homeostatic proliferation and sustain the HIV reservoir by occasional expansion and contraction of individual CD4 T-cell clones^19–21^. Specifically, increased expression of co-inhibitory receptors (PD-1, LAG-3, and TIGIT) that can drive cellular quiescence upon receptor engagement and inhibit HIV replication, have been shown to contribute to the magnitude of the HIV reservoir ^21–23^

In this study of two independent cohorts of HIV-infected cART-treated individuals, we identified INRs with high serum levels of anti-inflammatory molecules (triggered by elevated FOXO3 expression, TGF-β associated genes), a senescent CD4 T cell profile with absence of homeostatic proliferation (and lack of CD4 recovery), and heightened frequencies of cells with inducible HIV RNA.

## Results

We used an unbiased approach that integrates data-sets from peripheral blood transcriptome, high density flow cytometry and plasma cytokine measures, to identify cellular and molecular drivers of HIV persistence and lack of CD4 T cell recovery (Supplementary Fig. 1a), in two independent cohorts of HIV-infected subjects (see Study Participants in Material and Methods; Table S1, Table S2). All subjects were under cART for at least three years and maintained consistent CD4 counts four years prior to sample collection (Supplementary Fig. 1b). The Cleveland Immune Failure cohort (CLIF) was composed of 61 subjects (17 immune responders – IRs – >500 CD4 T-cells/mm^3^ and 44 immune nonresponders – INRs – <350 CD4 T-cells/mm^3^); whereas, the Study of the Consequences of the Protease Inhibitor Era cohort (SCOPE) included 41 subjects (20 IRs with >500 CD4 T-cells/mm^3^ and 21 INRs with <350 CD4 T-cells/mm^3^).

### Transcriptional profiling reveals systemic senescence as a driver of poor immune reconstitution

Exploratory analyses of whole blood transcriptomic data (CLIF cohort subjects; using unsupervised clustering described in the methods) (Fig. 1a and Supplementary Fig. 1c) identified three groups of subjects with unique transcriptional profiles: IRs and two distinct INR groups, “INR-A” and “INR-B”.

**Figure 1.**
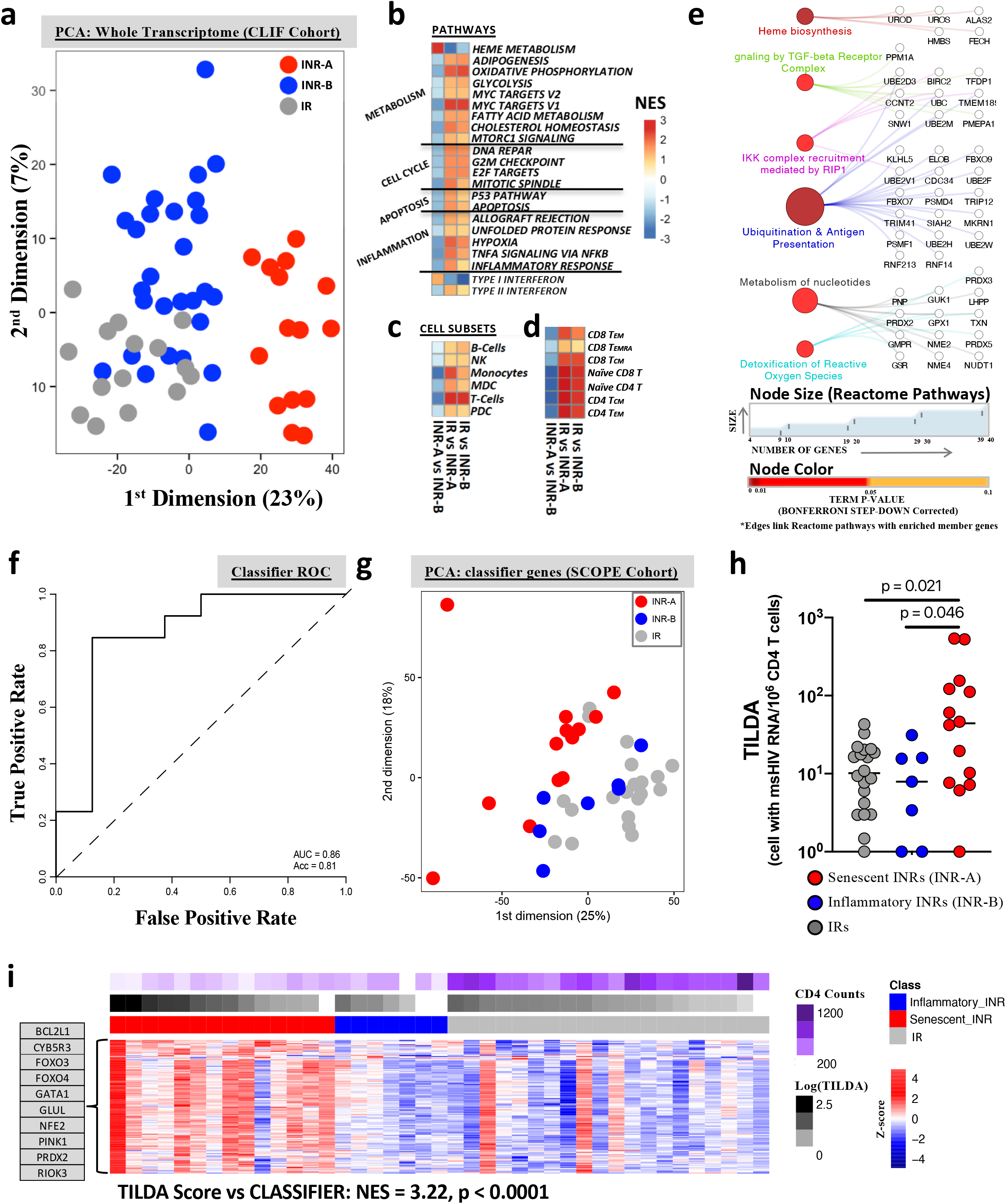
Whole blood transcriptional profiling identifies a 352-geneset signature that discriminates ART-treated Senescent-INRs and is associated with high inducible HIV. **a**, Multi-dimensional scaling (MDS) was used to reduce the Euclidean distance between whole blood samples of the CLIF cohort (n=61) into two dimensions that summarize the largest transcriptomic variation in the dataset. These two dimensions were used to represent the differences between samples (circles) in the scatter plot. Three groups were identified: IR (Grey; n=17), INR-A (Red; n=14), and INR-B (Blue, n=20). The first dimension of the plot, representing 23% of the transcriptomic variance between samples, depicts the observed transcriptomic difference between the three groups of subjects. **b-d**, Heatmaps illustrating the normalized enrichment score (NES; red (high) to blue (low) scale shown on the heatmap) of the top genesets (GSEA p-value ≤ 0.05) MSigDB: Hallmark^25^ (Table S6) + Interferome TypeI/II^76^ Pathways (**b**) Major immune cell Subsets^31^ (Table S7) (**c**) and T cell subsets^32^ (Table S8) (**d**) between *IR vs INR-A, IR vs INR-B and INR-A vs INR-B* subjects. Each row depicts a genesets, and columns represent the contrast between subject groups. **e**, Network highlighting the top biological functions associated with genes upregulated in INR-A subjects (vs IRs and INR-Bs; FDR < 0.05). As indicated, node color (dark red to orange) highlights the p-value resulting from over representation analyses, while the node size represents the number of genes per gene-set. Member genes are represented as white circles and connected to member geneset by grey edges (see Table S10 for details). Reactome database was used to annotate the biological functions and the network was plotted using the ClueGo plug-in within cytoscape. **f**, Receiver operating characteristics (ROC) curve for the genes as predictors of INR groups. X- and Y- axes represent the False and True positive rates of prediction on the SCOPE validation dataset. A 352 gene-based classifier (see Table S11 for list of genes) trained on the CLIF cohort segregates INR-A and INR-B subjects in the SCOPE cohort with an accuracy of 81%. The classifier which was tested across different microarray platforms confirmed the heterogeneity of ART-treated INR subjects in these two independent cohorts (see methods for details on approach used to build the classifier). **g**, Multi-dimensional Scaling was used to summarize the variation of the 352 genes of the classifier among subjects of the SCOPE cohort (n=41). Senescent-INR (Red), and INR-B (Blue) were identified along the first dimension of the MDS plot. **h**, Jitter plot illustrates significantly (Wilcoxon rank test p-values shown on plot) higher levels of inducible HIV (measured by TILDA^70^) in Senescent-INRs compared to IRs and INR-Bs. **i**, Heatmap illustrating the leading edge from the gene-based classifier that predict inducible HIV among subjects of the SCOPE cohort (n = 41; linear regression: classifier genes ~ inducible HIV, GSEA: NES=3.22, p<0.0001). Rows represent the z-score normalized genes in the leading edge and columns represent samples of the SCOPE cohort: IR (Grey), Senescent-INR (Red), and INR-B (Blue). As indicated in the legend, each row is z-score normalized. The magnitude of CD4 counts (purple) and the size of inducible HIV (black) are plotted as annotations at the top of the heatmap, as indicated. Selected genes are highlighted in boxes to the left of the heatmap (See Table S12 for full leading-edge gene list).

INR-As exhibited the highest transcriptomic variation from IRs with approximately 3000 differentially expressed genes (DEGs). In contrast, INR-Bs were proximal to IRs with <400 DEGs (Table S3-5). Age, years on cART, CD4 counts, CD4 nadir, markers of gut barrier dysfunction (sCD14), and inflammation (IL-6, IP10) – all known predictors of morbidity and mortality in INRs^24^ – failed to distinguish the two INR groups (Supplementary Fig. 1d-p). Interestingly, DEGs specific to INR-As were mostly down-regulated (72% and 66% when contrasted against IRs and INR-Bs, respectively), suggesting a quiescent transcriptional state within these subjects (Supplementary Fig. 2a).

Comprehensive pathway analyses using MSigDb’s Hallmark module^25–27^ (Fig. 1b, Table S6) revealed that several pathways, reflective of significantly elevated CD4 numbers, were upregulated in IRs when compared to both INR-As and INR-Bs. Notably, when comparing INR-As and INR-Bs (where CD4 numbers are comparable), INR-As showed significantly decreased expression of inflammation, cell cycling, apoptosis, and cellular metabolism genesets, indicative of a senescent state^28,29^. INR-As had reduced expression of genes regulated by Myc, a primordial TF that regulates pathways involved in proliferation and metabolism (glycolysis, oxidative phosphorylation and fatty acid metabolism) of activated T cells^30^.

Cell subset deconvolution (using gene signatures from Nakaya et al^31^) revealed a significant down-regulation in INR-As of genesets specific to all major innate and adaptive immune cell subsets (Fig. 1c, Table S7). Apoptosis, transcription/cell migration, and cellular/lipid metabolism (lipid storage, GTP metabolic process) were enriched in gene signatures down-regulated in T-cells, myeloid dendritic cells (mDCs) and monocytes, respectively (Supplementary Fig. 2b). Similarly, down-regulation of genesets (extracted from Novershtern et al^32^ – see methods for detail) specific to CD4 and CD8 T-cells naive, memory and effector cell subsets (Fig. 1d, Table S8). Altogether, these data provide supporting evidence for decreased global transcription and systemic senescence in immune cells from INR-As, when compared to INR-Bs or IRs.

### Transcriptional profiles downstream of FOXO3 and SMAD2/3 signaling are enriched in senescent INRs

To identify mechanisms underlying poor immune reconstitution observed in INR-As, we first identified TFs that were specifically enriched in INR-As (p < 0.05, Supplementary Fig. 2c, Table S9). These TFs included IRF3 (driver of type I IFN production^33^), FOXO3 (transcriptional repressor^34^), SMAD2 (TGF-β signaling^35^) and CCNT2 (Negative regulator of HIV Tat protein; i.e. driver of HIV latency induction^36^) (Supplementary Fig. 2c). Next, we used the Reactome pathway database (using ClueGO, FDR <0.05)^37,38^ to map the functions of upregulated genes in the INR-As (vs IRs and INR-Bs; FDR < 0.05, Table S10). We observed the enrichment of cellular processes downstream of the abovementioned TFs including heme metabolism, TGF-β signaling, IRF3 activation, reactive oxygen species (ROS) production and inhibition of NF-κB activation (Fig. 1e) in these subjects. Specifically, features of senescence regulated by FOXO3 including SOD/CAT^39–41^ (driver of ROS production) (Fig 1e); and FOXO4/TP53^42^ driven anti-apoptotic genes (BCL2L1; FDR < 0.01)^43–45^ were upregulated in INR-As. Interestingly, we observed the enrichment of key senescence associated genes which are known targets of SMAD2/3 including ‘Inducer of Promyelocytic Leukemia’ (PML; conductor of TGF-β signaling via SMAD2/3, and regulator of HIV latency^46,47^), WEE1^48^ (cell cycle regulator) and GLUL^49,50^ (metabolic regulator) (Supplementary Fig. 2d). These senescent features of INR-As were further confirmed by the downregulation MYC target genes (known to control ribosomal biosynthesis and translation)^51^, genes of the electron transport chain, and genes regulating major metabolic pathways (including glycolysis, oxidative phosphorylation and fatty acid metabolism) (Supplementary Fig. 2d, Fig. 1c).

The senescent/anti-inflammatory profile of INR-As contrasted the pro-inflammatory profile of INR-Bs where the enrichment of inflammatory pathways downstream of TLR/IL1, cell cycling, and apoptosis/pyroptosis (Supplementary Fig. 2d) was observed. Increased expression of several members of the pro-inflammatory NF-κB family of TFs (NFKB1, NFKB2 and RELB) (Supplementary Fig. 2d), upregulation of NF-κB target genes including chemokines (e.g. CCL2 and CCL17)^52^, genes of the inflammasome complex (NLRP3, IL1B and IL18)^53^ and effector genes driving apoptosis/pyroptosis (e.g. CASP4, DIABLO, CASP2 and CASP3)^54^ were characteristics of this group (Supplementary Fig. 2d).

Overall, our data suggest that a signaling cascade downstream of TGF-β signaling (SMAD2/3; PML) which culminates by the upregulation in the INR-As of FOXO3/4 driven senescence downstream of TGF-β and characterized by impaired cell metabolism and cell cycle arrest. The expression of the above listed transcriptional repressors in INR-As could further explain the inhibition of cell metabolism, cell cycle and inflammation all associated to cell senescence^55^. INR-As will henceforth be referred to as “Senescent-INRs”.

### A gene-based classifier confirms the generalizability of the Senescent-INR phenotype in HIV-infected subjects

Our next objective focused on demonstrating that other cohorts of HIV infected cART treated subjects also include senescent INRs. We built and validated a gene-based nearest shrunken centroid classifier (see methods) to identify Senescent-INRs in an independent cohort (SCOPE) of cART-treated HIV infected subjects. The classifier, trained on the CLIF cohort, consisted of 352 features (genes, Table S11) and provided a misclassification error rate of 0.26 (Supplementary Fig. 3a). Using the unsupervised approach (initially applied in the CLIF cohort), we confirmed two distinct INR groups in the SCOPE cohort (Supplementary Fig. 3b). The 352-gene classifier segregated these two INR groups in the SCOPE cohort with an accuracy of 81% (Fig. 1f, Supplementary Fig. 3c), thus validating the generalizability and reproducibility of two INR groups (PCA representation of the 352-gene classifier discriminates senescent INRs in Fig. 1g) and highlighting the potential use of this classifier to distinguish Senescent-INRs in clinical settings. This classifier (Table S11) included genes associated with the induction of a senescent profile (FOXO3, FOXO4, and TGFBR2)^43,56^, genes indicative of senescence biology (RIOK3, IRF3, BCL2L1)^57,58^ and regulators of mitochondrial activity (NDUFS3, CYB5R3, ATP5G2) Several transporters of macromolecules: SLC6A8 (Sodium- and chloride-dependent creatine transporter 1^60^), SLC48A1 (Heme transporter^61^), SLC4A1 (Band 3 anion transporter^62^), SLC25A23 (Mitochondrial Calcium carrier^63^) and SLC38A5 (glucose^64^) were also upregulated in Senescent-INRs and were features of the classifier (Table S11). These results higjlight several molecular pathways specific to cellular senescence^65–68^ and recapitulate the pathways (senescence, metabolism, and cell cycle) that discriminate between the two INR groups. These data counter-intuitively suggest that failure to increase CD4 T-cell numbers in senescent INRs to levels observed in IRs could be due to senescence and not to previously described pro-inflammatory cytokines and pathways^12^.

### The classifier gene set highlights the role of senescence in driving the magnitude of the inducible HIV reservoir

Although poor immune reconstitution has been associated with HIV persistence^69^, the cellular effectors and molecular mechanisms driving this association remain to be identified. To assess the role of senescence in HIV persistence, we measured frequencies of CD4 T-cells with inducible multi-spliced HIV RNA (“inducible HIV”) on all subjects of the SCOPE cohort using the Tat/rev Induced Limiting Dilution Assay (TILDA)^70^. Our data show that the Senescent-INRs have significantly higher inducible HIV when compared to IRs (~4.3-fold increase in median values, p<0.021) and INR-Bs (~5.5-fold increase in median values, p = 0.046) (Fig. 1h). As might be expected, inducible HIV levels were negatively correlated with CD4 counts, selectively in Senescent-INRs (rho = – 0.52; p < 0.05) while this correlation was not significant in inflammatory INRs and in IRs. In addition, SCOPE cohort subjects with the highest levels of inducible HIV showed a significant enrichment in (i.e. could be predicted by) the classifier geneset (Fig. 1i). Comprehensive analyses of the whole transcriptome confirmed that genesets characteristic of Senescent-INRs (i.e. cell cycle arrest and ROS production) were associated with higher inducible HIV (Table S12).

To better characterize the impact of senescence on intrinsic mechanisms of anti-viral activity, we studied the interplay of three master-regulators (IRF3: Type I IFN response^33^, IRF7: anti-viral response^71^, FOXO3: Senescence^42^) that showed significant alterations in senescent INRs (Fig 2). Increased expression of IRF3 (Fig. 2a) in Senescent-INRs was strongly associated with higher levels of FOXO3 (Fig. 2b), and contrasted the significantly reduced expression of IRF7 (Fig. 2c). These findings are in line with previous reports where a negative feedback loop between FOXO3 and IRF7^72^ has been ascertained, and suggest that Senescent-INRs could exhibit heightened inducible HIV when compared to IRs and INR-Bs as a consequence of the FOXO3 driven downregulation of innate antiviral immunity. Indeed, restriction factor genes known to curb HIV life cycle and known tobe transcriptional targets of IRF ^71,73^ were reduced (examples include SAMHD1, BST2, APOBEC3G and MX2) in subjects with the lowest inducible HIV (Fig. 2d, Table S12). In contrast, ChIP-seq validated targets of FOXO3/IRF3^74^,^75^ (Fig. 2d, Table S12) and type I IFN regulated genes^76^ (Table S12) that were correlated with higher inducible HIV included genes involved in promoting HIV replication (AFF1^77^, MT2A^78^, DARC^79^), induction of latency (BRD4^80^, GMPR^81^) and persistence of HIV infected cells (BCL2L1^82^). These data show that lower levels of IRF7 as a consequence of higher IRF3/FOXO3 lead to dysregulated antiviral innate immunity and heightened inducible HIV in the senescent INRs.

**Figure 2.**
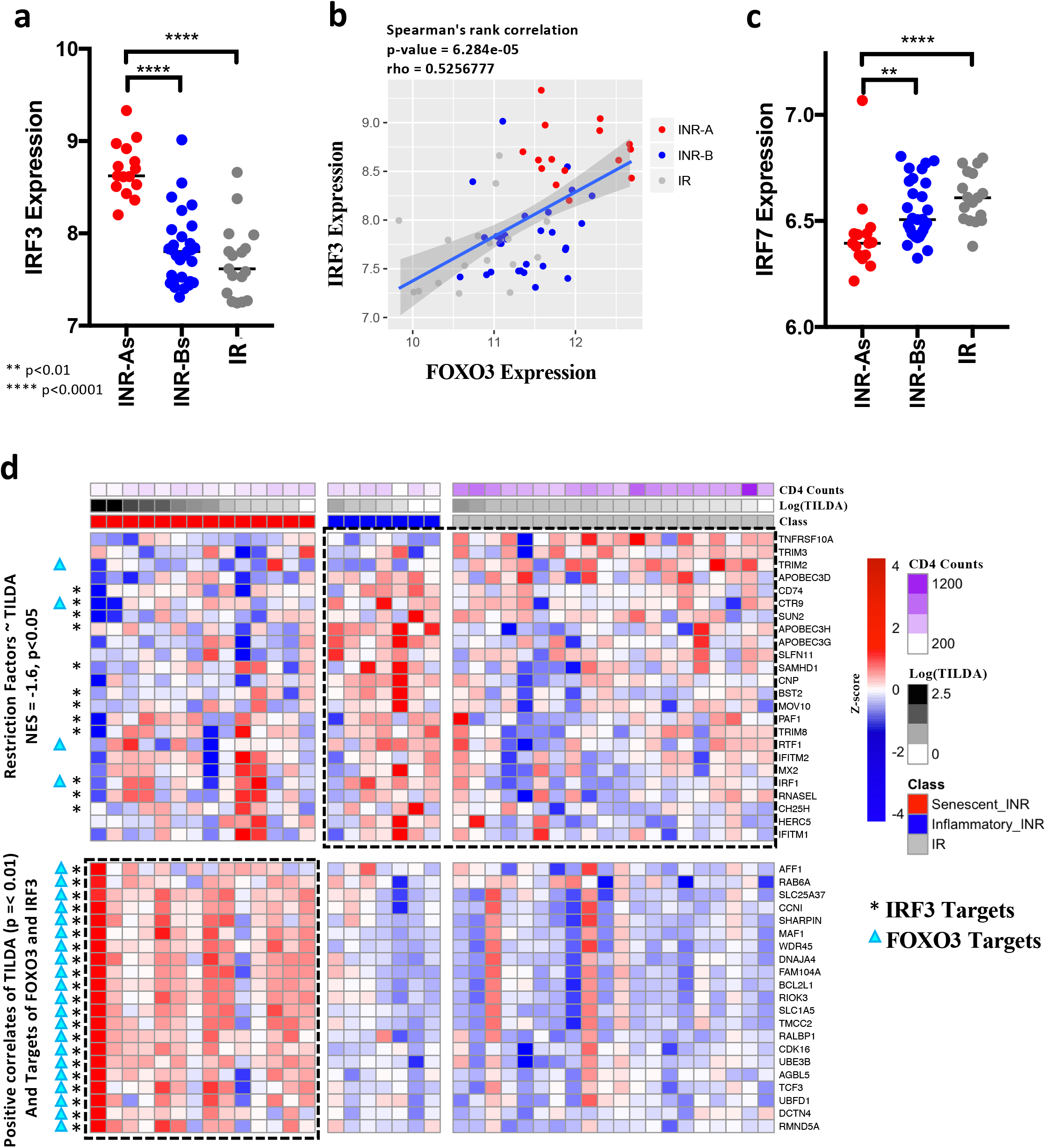
Altered interferon signaling drives inducible HIV in Senescent INRs. **a-b**, Jitter plot illustrating significantly elevated levels of IRF3 expression in Senescent-INR subjects (a), scatter plot showing a significant positive correlation between FOXO3 and IRF3 expression (b) and diminished levels of IRF7 expression in Senescent-INR subjects (c) of the CLIF cohort. A Wilcoxon rank sum test was used to assess significant differences between groups (**p < 0.01, ****p<0.0001), and a spearman correlation test was used to assess significance of association (p-value and rho indicated on plot). **d**, Heatmap illustrating the dichotomy of interferon signaling driving the HIV reservoir size among subjects of the SCOPE cohort (Top block: linear regression: Restriction Factors ~ inducible HIV, GSEA: NES= – 1.6, p<0.05, Bottom block: positive correlates of inducible HIV that are targets of FOXO3 and IRF3). Rows represent the z-score normalized genes in the leading edge and columns represent samples of the SCOPE cohort: IR (Grey), Senescent-INR (Red), and INR-B (Blue). As indicated in the legend, each row is z-score normalized. The magnitude of CD4 counts (purple) and the size of inducible HIV (black) are plotted as annotations at the top of the heatmap, as indicated. See Table S12 for full gene-lists.

### Signalling via TGF-β is a hallmark of senescent INRs and directly drives HIV latency in memory CD4 T cells in vitro

Transcriptional profiling identified the TGF-β pathway (PML, SMAD2/3; Fig. 1e, Supplementary Fig. 2c,d) as a driver of cellular senescence in our cohort. T-regulatory cells (Tregs) that express Forkhead box P3 (FOXP3) are a primary source of TGF-β and are critical in maintaining immune homeostasis^83,84^. To ascertain the role of Tregs in driving senescence and HIV persistence, we developed a high-dimensional flow cytometry panel (Table S13) to quantify master regulators of Treg function (SATB1 and FOXP3^85^), to discriminate differentiated Tregs (CD45RA, CD49B and CD39^86^), to assess proliferation (Ki67) and to measure the capacity to activate TGF-β by expression of GARP and/or LAP (two molecules involved in promoting latent (LAP) or activated (GARP) forms of TGF-β^87,88^). Density projection of CD4 T cells expressing these markers on a two-dimensional UMAP (https://arxiv.org/abs/1802.03426) revealed distinct distribution of cells within Senescent-INRs (Fig. 3a). Next, we used unbiased clustering^89^ (Fig. 3a) to identify three clusters of CD4 T cells (Fig. 3b, Table S14) that were enriched in senescent INRs (vs IRs) and expressed markers characteristic of TGF-β producing Tregs (FOXP3, CD25, lowCD127, LAP and GARP) (Fig. 3c). The most abundant of these clusters (i.e. Cluster 7; 0.81-5.34% of CD4 T cells) also showed an effector Treg phenotype (high CD39, low CD45RA) and low levels of markers that define IL-10 producing Tregs (Tr1) (i.e. CD49B, LAG3^90^) (Fig. 3c).

**Figure 3.**
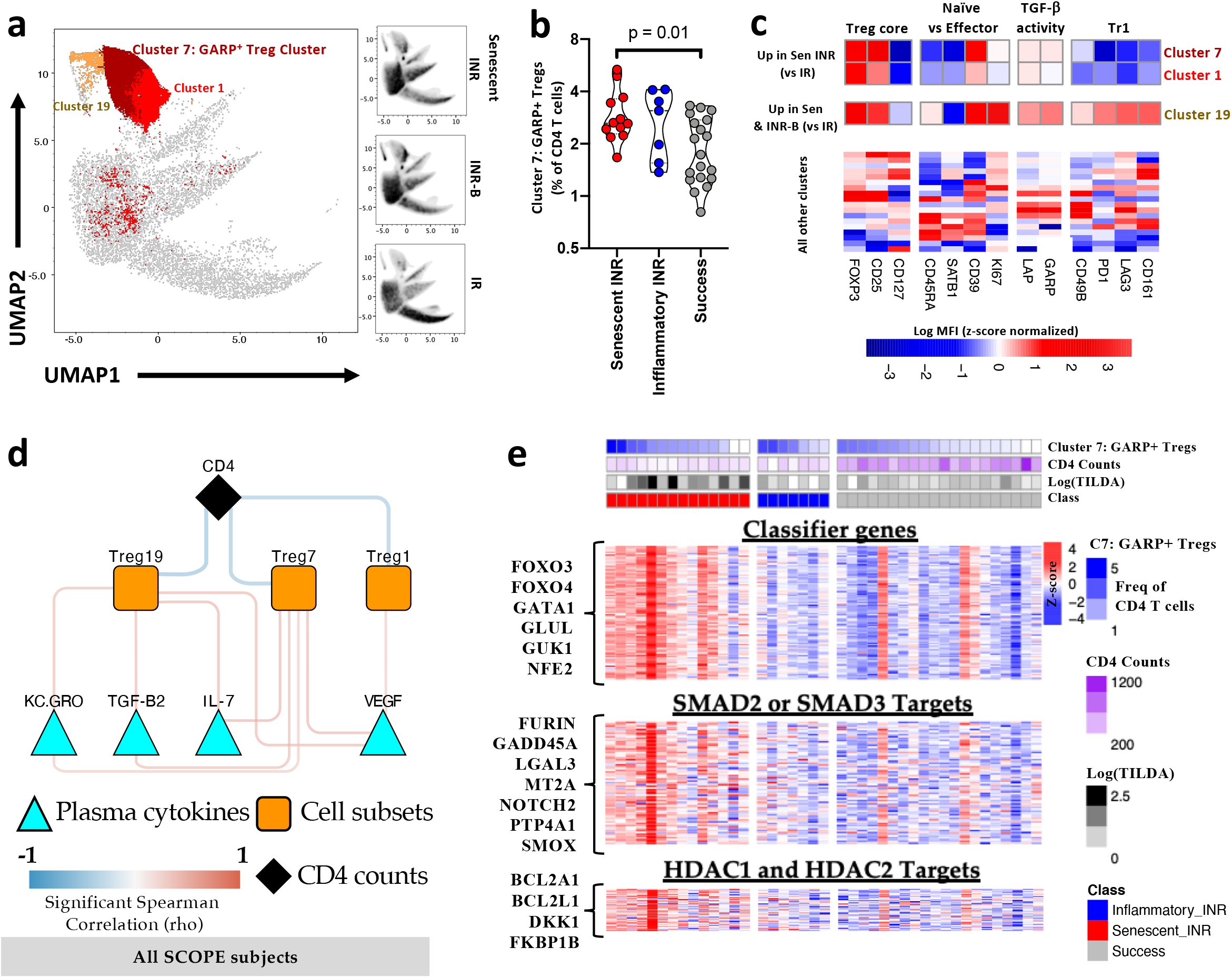
TGF-β signaling cascade exemplified by an increase in SMAD2/3 targets and an increase in TGF-β producing Tregs is associated with an increase in inducible HIV and drives latent HIV in vitro. **a**, Treg panel clusters determined using the PhenoGraph method (see Table S14) and visualized using the Uniform Manifold Approximation and Projection (UMAP) analysis were used to represent the distribution of Treg phenotypes in total CD4 T cells (using highdimensional flow cytometry) in IR, Senescent-INR, and INR-B subjects of the SCOPE cohort. FOXP3 expressing clusters that were abundant in Senescent-INR subjects (vs IRs) are highlighted on the UMAP plot (See Table S14 for details). **b**, Violin plot illustrates significantly higher frequencies of Treg cluster 7 (in total CD4 T-cells) in Senescent INR subjects. Wilcoxon rank test was used to assess significance and p-values are indicated on the plot. **c**, Heatmap highlighting the differences in markers between clusters. FOXP3+ clusters that were abundant in senescent INRs (i.e. clusters 1,7, and 19) are highlighted. Each column is z-score normalized value of the mean fluorescence intensity (raw intensities for every cluster can be found in Table S14). **d**, Network highlighting a positive correlation between FOXP3 expressing Treg clusters that are abundant in Senescent INRs (Clusters 1, 7 and 19) with plasma cytokine levels (of cytokines in cytokine cluster 3; See Supplementary Fig. 4b) measured on the same subjects. Triangular blue nodes depict plasma cytokines IL-7, TGF-β2, KC.GRO and VEGF; red edges highlight a positive correlation between those cytokines and the Treg clusters. Orange squares reflect high-density flow cytometry clusters, light blue triangles reflect plasma cytokine levels, red edges indicate significant positive correlation. A Spearman correlation test was used to assess significance (p-value <0.05) across IR, Senescent-INR, and INR-B subjects of the SCOPE cohort (Table S16 lists the details of correlations between plasma cytokine cluster 3 members and Treg subsets). **e**, Heatmap highlighting overlapping leading-edge genes from the association of inducible HIV and frequencies of Treg cluster 7 with the 352 gene-based classifier genes (top block), SMAD2/3 targets (middle block) and HDAC1/2 targets (bottom block). Rows represent features and columns represent samples of the SCOPE cohort: IR (Grey), INR-A (Red), and INR-B (Blue). The gene-expression per row was centered at zero and to a standard deviation of 1 (z-score). As indicated in the figure, a red-white-blue gradient is used to depict the relative expression of the features. The magnitude of CD4 counts (purple), frequencies of Treg Cluster7 (GARP+, dark blue) and size of HIV inducible reservoir (black) are plotted as annotations at the top of the heatmap. GSEA was used to assess the association between the features and inducible HIV/Treg cluster 7 frequencies (linear regression, p<0.05). Genes that showed a significant overlap (GSEA vs inducible HIV and Treg cluster 7 frequencies; significance (p-value <0.05 – assessed by Fischer exact test) amongst the leading-edge gene list/pathway are represented on the heatmap (See Table S17 for all the details). **f-g** The LARA in vitro model^112^ was used to characterize the impact of dose-dependent increase in TGF-β on the establishment of HIV latency in memory CD4+ T cells. Increasing concentrations of TGF-β (0.2-20ng/mL) during the latency phase of the assay led to heightened frequencies of CD4 T cells with integrated HIV DNA **(f)** and resulted in significantly higher frequencies of p24+ CD4 T cells upon reactivation with anti-CD3/28 antibodies **(g)**. Spearman’s test was used to assess significance for correlation in (f) (p-value and rho as indicated on graph) and Wilcoxon rank-sum test was used to assess significance between timepoints in (g) (p-value as indicated on graph).

We assessed the systemic impact of these Treg clusters by studying their association with groups of cytokines that define the host plasma milieu. To this end, unsupervised clustering of the 43 different plasma cytokines using independent methods (k-means, and hierarchical clustering^91^) was performed and revealed four clusters of plasma cytokines across all subjects of the SCOPE cohort (Table S14, Supplementary Fig. 4a,b,c). Of these clusters, the overall centroid score of cytokine “Cluster 3” showed significant association with the 352 classifier genes in Senescent-INRs (NES = 2.9, FDR = 0, Supplementary Fig. 4d, Table S15). No significant correlation between cytokine Cluster 3 centroid score and the classifier genes was observed INR-Bs or IRs (Table S15). Cluster 3 included several anti-inflammatory cytokines: TGF-β1, TGF-β2 (triggers of the TGF-β pathway^92^), IL13 (known to inhibit inflammatory cytokine production^93^), KC/GRO (CXCL1) (another anti-inflammatory cytokine^94^), VEGF (the inhibitor of apoptosis^95^) as well as growth and homeostatic cytokines (IL3, IL7)^96,97^ (Supplementary Fig. 4c). Importantly, our data showed that frequencies of GARP+ Tregs (Treg cluster 7) were univariately correlated with members of cytokine cluster 3 (Fig. 3d, including TGF-β2 and IL-7; Table S16), and were associated with an enrichment in the classifier genes that were predictive of inducible HIV (significantly overlapped leading edge genes represented in Fig. 3e, Table S17).

The downstream impact of heightened frequencies of TGF-β secreting Tregs on HIV latency was confirmed by the observation that ChIP-Seq validated SMAD2/3 targets^98^ and HDAC1/2 targets (induced after siRNA knockdown)^99^ were more abundant in subjects with the highest inducible HIV and frequencies of GARP+ Tregs (Fig. 3e). These genesets were characterized by enhanced expression of genes that activate latent TGF-β (i.e. FURIN^100^), restrict T cell differentiation (LGAL3; inducer of TGF-β driven activation of β-catenin^101,102^) and reduce cell cycling (i.e. GADD45A^103^). The co-operation between HDAC1/2 targets (i.e. BCL2L1, FKBP1B) and SMAD2/3 can restrict chromatin accessibility and induce cell quiescence (Fig. 3e).

### Microbial metabolites drive Treg differentiation and TGF-β production in Senescent-INRs

Bacterial translocation is a known feature of HIV pathogenesis in ART-treated and untreated HIV-infected subjects^104^. INRs of the CLIF cohort (Supplementary Fig. 1m) exhibited significantly higher levels of sCD14 (marker of disease progression and leaky gut^105^) compared to IRs. However, Senescent-INRs and INR-Bs showed comparable sCD14 levels (Supplementary Fig. 1m, p=0.567). We therefore monitored for qualitative changes in the microbiome (prevalence of different bacterial quasispecies), which could be associated to the different transcriptomes, cytokine environments and T-cell subset distributions that distinguished the two groups of INRs. Of note, several bacterial metabolites produced by commensal bacteria (i.e. acetates, butyrates, and propionate) have been shown to regulate Treg differentiation^106,107^ and T-cell homeostasis. We examined the host microbiome of INRs using PathSeq^108,109^ to obtain an overall quantitative and qualitative measure of the viral, bacterial, fungal, parasite and helminth burden in subjects of the SCOPE cohort. We performed mass spectrometry (MS) to measure plasma metabolite levels and to identify bacterial metabolites that would be associated (*ex vivo)* or could trigger *(in vitro)* Treg differentiation, TGF-β production or senescence, and monitor their association with Treg frequencies and levels of senescent cytokines.

Using PathSeq, we first observed higher Shannon diversity (Supplementary Fig. 5a) and increased abundance of the Firmicute phylum in the Senescent-INRs (Fig. 4a, p=0.05). All other bacterial phyla that were detectable by Path-Seq were not significantly different in abundance between Senescent and inflammatory INRs (Fig. 4a). Furthermore, we observed that overall species diversity of the Firmicute phylum (assessed by the Bray-Curtis dissimilarity statistics^110^) significantly distinguished the two groups of INRs (Supplementary Fig. 5b). To granularly assess the differences between the Senescent-INRs and inflammatory INRs, we generated a NCBI Taxonomy tree (newick tree files created using the phyloT – http://phylot.biobyte.de – and visualized using the iTOL – http://itol.embl.de – web-interfaces, respectively) of all bacterial genera with median abundance > 0 TPM in either of the three groups in the SCOPE cohort. We observed that the Senescent-INRs (vs inflammatory INRs) were significantly enriched in genera from all major phyla. These included general like *Moraxella/Bradyrhizobium* (Proteobacteria), and Actinobacteria like *Kytococcus/Mycobaacterium* (Fig. 4b). More importantly, we observed that the abundance of *Lactobacillus* (an anti-inflammatory genus and a member of the Firmicute phylum – i.e. phylum higher in Senescent-INRs) and the Lactobacillaceae family were significantly correlated with inducible HIV levels (Fig. 4b) (Supplementary Fig. 5c). Altogether, these data support microbiome changes specific to Senescent-INRs; and pinpoint the presence of Firmicute species in plasma that are associated with magnitude of the HIV reservoir in Senescent-INRs.

**Figure 4.**
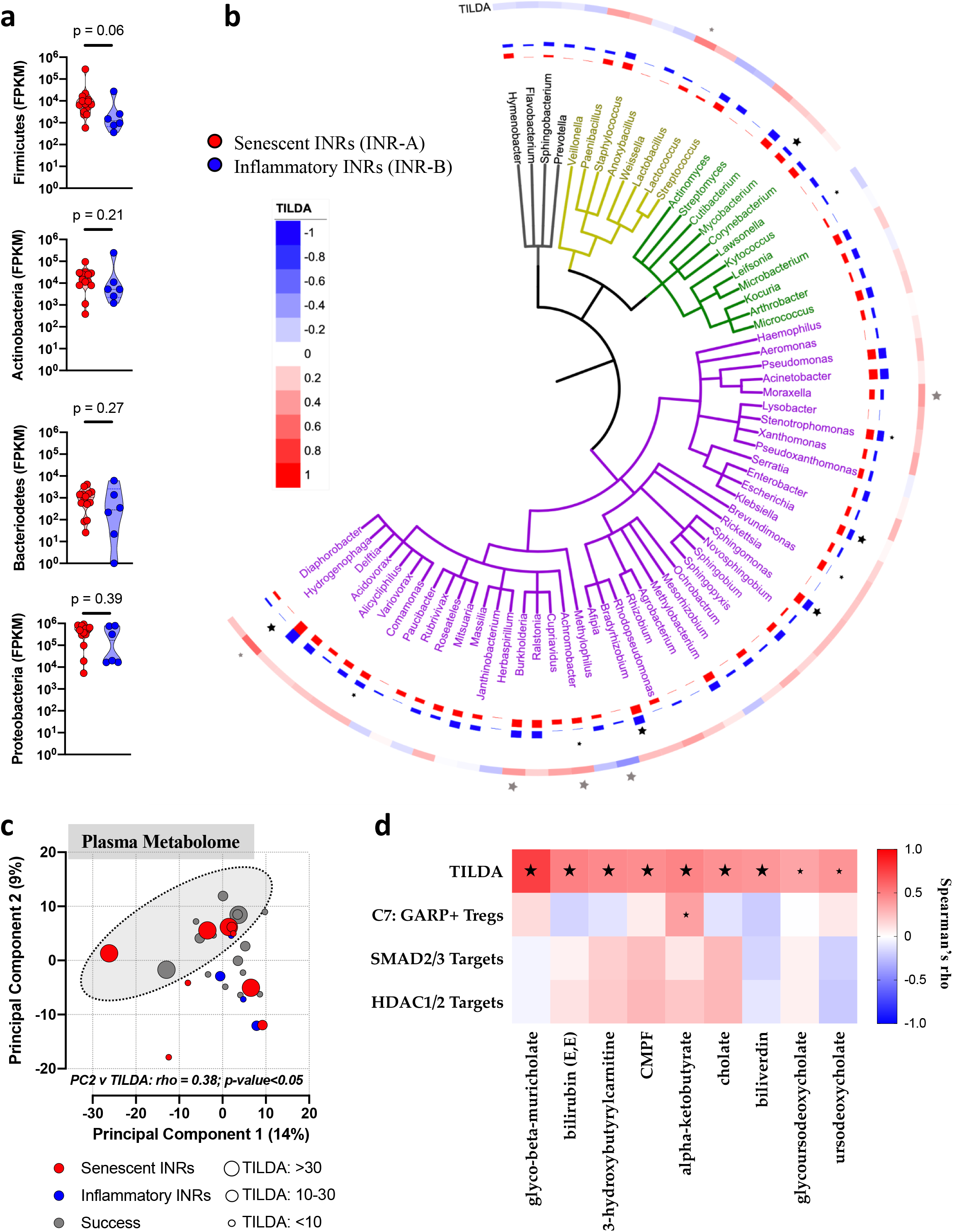
Microbial metabolites drive Treg differentiation and TGF-β production in Senescent INRs. **a**, Box plots illustrating levels of bacterial phylum Firmicutes are heightened in Senescent-INRs (p=0.06) of the SCOPE cohort. Subject groups are plotted along the X-axis: INR-A (Red), and INR-B (Blue) and log_10_ of relative Firmicute abundance based on PathSeq plotted along the Y-axis. A Wilcoxon tank sum test was used to assess significance. **b**, Taxonomy tree of all genera with abundance >0 TPM in all subgroups (IRs, Senescent and Inflammatory INRs) of the SCOPE cohort. Each genus label and tree leaf is colored based on bacterial phyla annotation. Bar plots in the concentric circles represent log(TPM+1) value for each genus in Senescent or Inflammatory INRs form the SCOPE cohort. Stars beside the bars highlight significant differences between Senescent or Inflammatory INRs (P<0.05: large start, P<0.1: small star). Heatmap strip in the concentric circle shows the correlation value (Spearman’s rho) of each genus with inducible HIV. Stars beside the heatmap strip highlight significant correlations (P<0.05: large star, P<0.1: small star). **c**, Principal Component Analysis of all plasma metabolome highlights a cluster of subjects with high TILDA levels and reveals a significant negative correlation between PC1 (x-axis) and the Classifier gene-set and a significant positive correlation between PC2 (y-axis) and TILDA. Spearman Correlation was used to assess significance, rho and p-values provided on the figure. **d**, Correlation matrix heatmap between abundance of plasma bile acid/microbial metabolites (plotted along the y-axis) and variables of interest including TILDA levels, GARP+ Tregs, SMAD2/3 and HDAC1/2 Targets (plotted along the y-axis). Analysis highlights α-ketobutyrate as the metabolite correlated with levels of TILDA and GARP+ Tregs. A spearman correlation was used to assess significance. Stars on the heatmap highlight significant correlations (P<0.05: large star, P<0.1: small star).

Bacterial translocation is known to have a major impact on host and systemic microbial metabolome which leads to alterations of innate immune responses. Hence, we assessed the drivers of variation in total plasma metabolite levels in the SCOPE cohort. The primary components of variation (Principal Component 1 and Principal Component 2; derived from PCA analyses of ~750 detectable metabolites) were associated with inducible HIV (PC2 vs TILDA p-value <0.05; Fig. 4c) and the classifier geneset (PC1 vs classifier genes that predict inducible HIV – p-value <0.05; Fig. 4c). The PCA analyses confirmed the role of the systemic metabolome in driving inducible HIV. Given the association of the higher *Lactobacilli* abundance with inducible HIV, we performed univariate analyses to identify plasma microbiome derived metabolites (i.e. short-chain fatty acids and bile acids) that correlated with inducible HIV levels, These metabolites included members of the liver-biliary axis (i.e. primary liver/bile metabolites like bilirubin, biliverdin, cholate and glycol-beta-muricholate; secondary liver/bile metabolites like ursodeoxycholate), carnitine derivatives and members of the hydroxybutyrate family (i.e. α-ketobutyrate and hydroxybutyryl carnitine) (Figure 4d). Several, but not all, of these metabolites were also associated with SMAD2/3 and HDAC1/2 target genesets, C7 GARP+ Tregs and inducible HIV (Fig. 4d). Importantly, this analysis showed that α-ketobutyrate was significantly correlated (p = 0.089) with C7 GARP+ Treg frequencies, emphasizing the association between Tregs, metabolome and inducible HIV. Together these data indicate that microbial metabolites and the microbiome could constitute integral components of the mechanisms that fuel HIV persistence.

### TGF-β production resulting from alpha-ketobutyrate driven Treg differentiation causes increased HIV latency in vitro

To establish a causal link between butyrate metabolites on GARP+LAP+ Treg differentiation, we used the approach described by Ohno and Rudensky^106,107^. Briefly, we assessed the impact of increasing concentrations of alpha-ketobutyrate on sorted naïve CD4 T-cells obtained from uninfected subjects in the presence of IL-2, anti-CD3/28 and/or TGF-β stimulation (Fig. 5a). TCR stimulation in the presence of TGF-β led to a profound increase in frequencies of FOXP3+ cells^111^. Addition of increasing concentrations of alpha-ketobutyrate in the absence of TGF-β preferentially led to the differentiation of naïve T-cells into GARP+FOXP3+ cells (Fig. 5a,b). Of note, TGF-β synergized with alpha-ketobutyrate to drive the differentiation of GARP+ Tregs (Supplementary Fig. 5d). This was supported by increased TGF-β1 cytokine secretion in the supernatant of naïve CD4 T-cell cultures stimulated with alpha-ketobutyrate (Fig. 5c). Furthermore, increasing concentrations of alpha-ketobutyrate led to significantly heightened levels of PD1 expressing CD4 T-cells (Fig. 5d). alpha-ketobutyrate induced GARP+ Treg (TGF-β producing) and PD1 + CD27+ phenotypes were also associated with loss of effector function – shown by reduced production of T-helper cytokines (i.e. IL17A, IFN-γ, IL9) (Supplementary Fig. 5f). Altogether, these data indicate that in addition to enhancing Treg differentiation, the abundance of hydroxybutyrates could drive the upregulation of TGF-β processing/suppressive activity of Tregs. The latter, as shown in Fig 3, are critical for the maintenance of the HIV reservoir.

**Figure 5.**
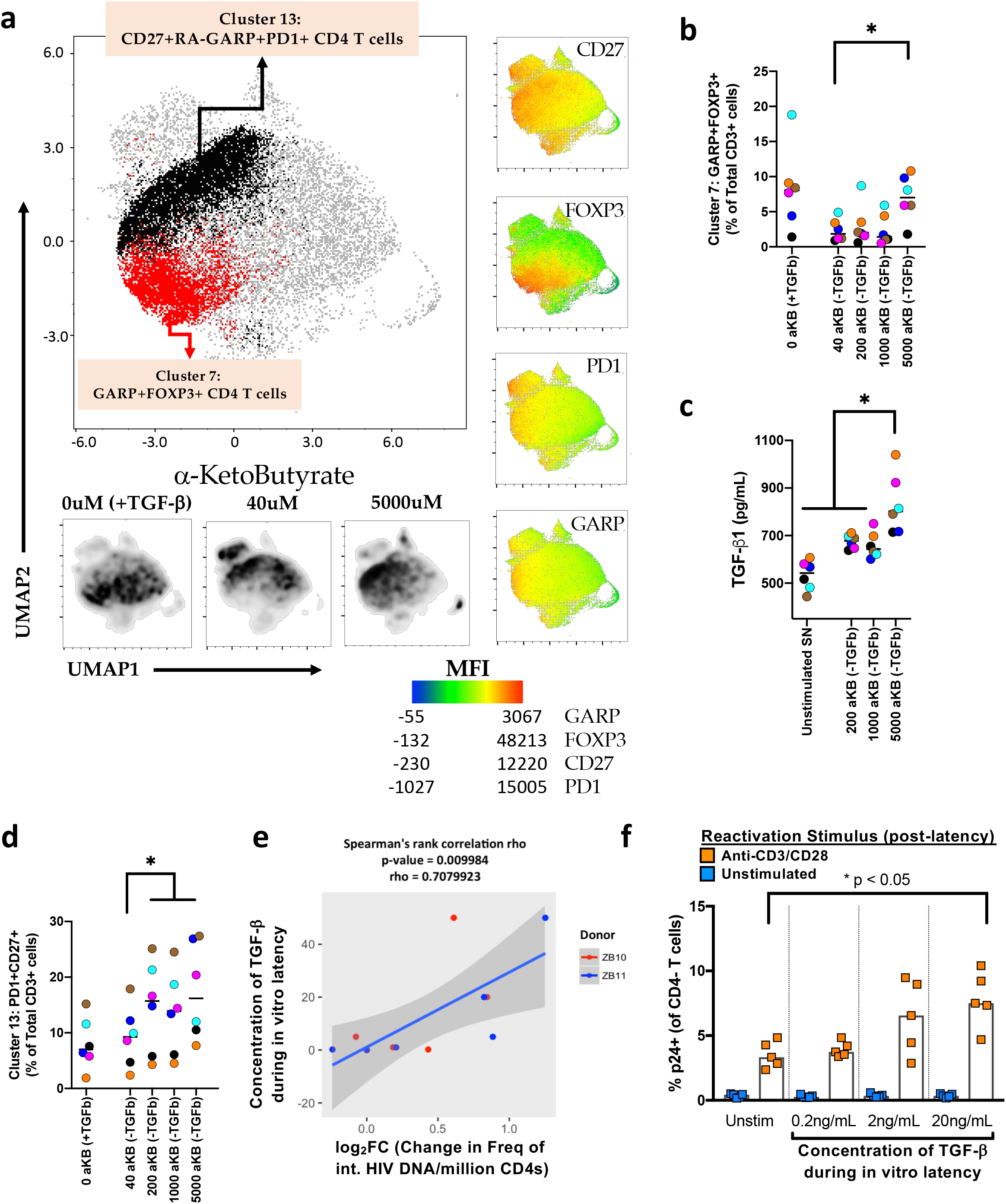
alpha-keto butyrate stimulation drives Treg differentiation and enhances TGF-β production. **a**, *in-vitro* experiment to assess the impact of increasing concentrations of alphaketobutyrate on sorted naïve CD4 T cells from healthy subjects in the presence of IL-2, anti-CD3/28 antibodies and/or TGF-β. Dimension Reduction of high density flow cytometry analysis highlights UMAP1 and UMAP2 on X- and Y-axes respectively and illustrates that stimulation in the presence of TGF-β led to a profound increase in frequencies of GARP+FOXP3+ cells (**a**, Cluster 7) and GARP+PD1+ cells (**a**, Cluster 13). **b, d** Jitter plots highlighting that increasing concentrations of alpha-ketobutyrate preferentially led to differentiation into GARP+FOXP3+ cells (**b**) and increased levels of PD1 expressing quiescent cells (**d**). GARP+FOXP3+ and PD1+CD27+ levels plotted along the Y-axis, while alpha-ketobutyrate increasing concentration levels are indicated on the X-axis. Conditions in the absence of TFG-b are designated with (-TGF-β). A Mann-Whitney U-test was used to assess significance across concentrations (* represents p-value <0.05 between groups). **c**, Plasma cytokine levels of TGF-β1 significantly increase with enhanced alphaketobutyrate stimulation. TGF-β1 levels plotted along the Y-axis, while alphaketobutyrate concentration levels are indicated on the X-axis. A Mann-Whitney U-test was used to assess significance across concentrations (* represents p-value <0.05 between groups). **e-f,** LARA assay in vitro model was used to characterize the impact of dosedependent increased in TGF-β on the establishment of HIV latency in memory CD4+ T cells. Increasing concentrations of TGF-β (0.2-20ng/mL) during LARA latency culture results in heightened frequencies of CD4 T cells with integrated HIV DNA **(e)** and results in augmented numbers of p24+ CD4 T cells triggered by reactivation using anti-CD3/28 antibodies stimulation **(f)**.

To provide evidence for a direct role of TGF-β in the induction of HIV latency, we developed an *in vitro* culture model^112^ where TGF-β was added to HIV-infected CD4 T-cells. Increased numbers of CD4 T-cells with integrated proviral DNA were observed after 14 days of culture with increasing concentrations (0 to 50 ng/mL) of TGF-β (r= 0.7; p = 0.009) (Fig. 5e), demonstrating that TGF-β contributes to heightened levels of non-replicative forms of HIV DNA and establishment of the HIV reservoir. We reversed latency in this system by stimulating CD4 T cells induced to become latent by addition of increased concentrations of TGF-β (0.2-20ng/mL) induced with immobilized CD3 and soluble CD28 specific antibodies. Our data show that frequencies of HIV p24+ cells were significantly higher in conditions where the highest concentrations of TGF-β (0.2-20ng/mL) were added (Fig. 5f). Altogether, these *in vitro/ex vivo* observations validate the critical role of TGF-β producing Tregs in driving the mechanistic establishment and/or maintenance of the HIV reservoir.

### Increased frequencies of senescent PD1+ TCMs with impaired mitochondrial function drive lower CD4 counts in the senescent INRs

Given that signaling via TGF-β/FOXO3 axis drives surface PD1 expression^113^, cell cycle arrest and impaired metabolism^114^ – we used highdimensional flow cytometry, transcriptomics and cytokine data to identify the cellular drivers of immune reconstitution or lack thereof; we hypothesized that the upregulation of protein levels of PD-1 and reactive oxygen species (ROS; measured by intracellular CellROX staining) in CD4 T-cells could be associated with lower CD4 counts^115^. Using the analytical strategy described in the above section, we identified a cluster of PD1^hi^ ROS^hi^ CD4 central memory T-cells (TCM, Cluster 9; Fig. 6a,b,c, Table S18) that was uniquely up-regulated in Senescent-INRs (i.e. not higher in INR-Bs vs IRs) when compared with IRs. Conversely, a cluster of PD1^hi^ ROS^lo^ CD4 effector memory T-cells (TEM, Cluster 17; Fig. 6a,b,c, Table S18) was uniquely upregulated INR-Bs. In line with previous studies^21,116^, these data confirm the negative impact of PD-1 expression on T-cell homeostasis and immune reconstitution.

A granular assessment of the plasma cytokine milieu revealed that levels of cytokines that drive effector differentiation (i.e. IL-15^112^) were directly correlated with increased frequencies of PD1^hi^ ROS^lo^ CD4 TEMs in INR-Bs (vs IRs) (p<0.05, Table S19). Whereas, increased frequencies of PD1^hi^ ROS^hi^ CD4 TCMs in the senescent INRs (vs IRs) were directly associated with higher plasma IFNα/IL-29 (type I and III IFN; two cytokines that drive PD1 expression^117^ and lower levels of fractalkine (a chemotactic factor downstream of IL-15 driven effector differentiation^118^) (Fig. 6d), indicating that unlike in the IRs, CD4 TCMs in senescent INRs express PD1, have impaired mitochondria and are stalled at the TCM stage. Evidence of impaired metabolism and cellular senescence in these PD1^hi^ ROS^hi^ TCMs was obtained by identifying metabolic and cell cycling genesets that associate with this cell subset (in senescent INRs vs IRs). Our data show that individuals with higher levels of PD1^hi^ROS^hi^ TCMs had poor mitochondrial metabolism profiles (higher ROS and lower oxidative phosphorylation) (Fig. 6e, Table S20), reduced cMyc activity (Fig. 6e, Table S20) and increased expression of genes that drive cellular senescence (Fig. 6f, Table S20). Here, expression of catalase (CAT), peroxidins (PRDXs) and cell cycle inhibitory genes (i.e. CDKN2D) was increased in senescent-INRs; while genes that regulate oxidative phosphorylation (NDUFs and COXs) and MYC target genes that drive transcription (RPSs)/translation (EIFs) were expressed at lower levels. Concurrently, frequencies of PD1^hi^ROS^hi^ TCMs were also associated with higher cellular senescence pathways downstream of impaired mitochondrial activity (i.e: oxidative stress induced senescence, SASP) (Fig. 6f, Table S20), and were observed to be the highest in subjects with higher inducible HIV (Fig. 6e, Table S20). Altogether, these results indicate that PD1 expressing TCMs with impaired mitochondrial metabolism are senescent cells lacking the capacity to cycle and to differentiate into effector cells, thereby driving low CD4 cellular counts.

### Integrated multi-omic model highlights cellular and molecular effectors of senescence as drivers of HIV persistence and lack of immune reconstitution

Using unbiased and holistic approaches, we highlighted gene expression profiles, cytokines and T-cell subsets that were independently associated with HIV persistence and lack of CD4 reconstitution in the Senescent INRs. To investigative the interplay of pathways driving these cellular and viral phenotypes and HIV persistence, we integrated multi-omic signatures above (Figures 2, 3 and 4, Table S21) across all subjects (n=41). Our data show that the levels of inducible HIV were positively correlated to senescence as characterized by impaired metabolism^115^ (increased ROS, decreased OX/PHOS), reduced transcription/translation (RPSs and EIFs) downstream of poor cMYC activity and TGF-β signaling (SMAD 2/3 target genes) (Fig. 6g, Supplementary Fig. 6). The induction of these cascades was consistent with higher expression of the transcriptional repressors that induced senescence (i.e. FOXO3, FOXO4)^42^, and with lower expression of master regulators of innate antiviral activity (i.e. IRF7 and restriction factors of viral replication) (Fig. 6, Supplementary Fig. 6).

**Figure 6.**
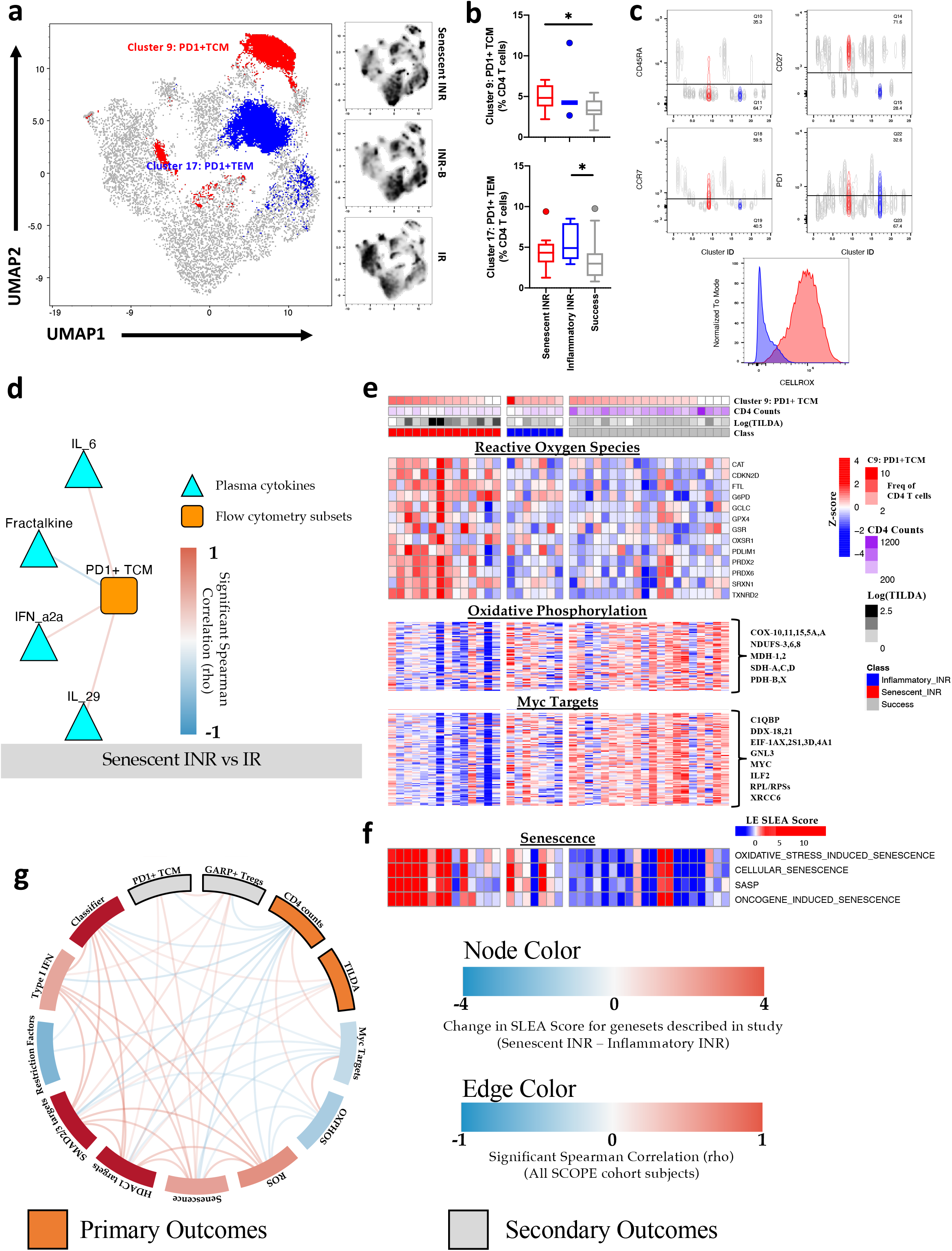
Heightened frequencies of PD1^hi^ROS^hi^ CD4 TCMs and lower CD4 counts in Senescent INRs are associated with poor mitochondrial metabolism and senescence. **a**, Clusters determined using the PhenoGraph method and visualized using the Uniform Manifold Approximation and Projection (UMAP) analysis were used to represent the distribution of CD4 T cells subsets, surface expression of PD-1 and markers of mitochondrial activity in CD4 T cells (using high-dimensional flow cytometry) in IR, Senescent-INR, and INR-B subjects of the SCOPE cohort (See Table S18 for details). Density Plots highlight the enrichment of a specific cluster “Cluster 9: PD1+TCM” in Senescent-INR subjects. **b**, Box plots illustrate unique and significantly higher frequencies of Cluster 9 (PD1+TCM in total CD4 T-cells) in Senescent INR subjects and Cluster 17 (PD1+TEM in total CD4 T cells) in the Inflammatory INR. A Wilcoxon rank test was used to assess significance (*p-value of <0.05) (Table S18). **c**, MFIs of cell surface (CD45RA, CD27, CCR7, PD1) per cluster are shown. Clusters 9 (red) and 17 (blue) are overlaid to highlight the heightened levels of CD27 and CCR7 in Cluster 9 (i.e. TCM), and low levels of CD27 and CCR7 in Cluster 17 (i.e. TEM). Both clusters showed relatively low levels of CD45RA and high levels of PD1. Black line across each plot discriminated traditionally gated negative population from the its positive counterpart. A head-to-head comparison between clusters 9 and 17 revealed a distinct upregulation in CellROX within cluster 9 (i.e. PD1+ TCMs that are increased in the senescent INRs) (Raw MFIs of all clusters are listed in Table S18). **d**, Correlation network highlighting significant associations between PD1+TCM cluster frequencies and plasma cytokine (positive: IL6, IL29, IFNa2a, negative: Fractalkine) in the pool of Senescent INRs and IRs (i.e. driven by lower CD4 counts in Senescent INRs vs IRs). Triangular blue nodes depict plasma cytokines; orange squares reflect high-density flow cytometry clusters; and red edges highlight a positive correlation between cytokines and cell clusters. A Spearman correlation test was used to assess significance (all edges have a significant p-value <0.05) across IR and Senescent-INR subjects of the SCOPE cohort (details of all correlations are listed in Table S19). **e**, Heatmap highlighting a positive association between Reactive Oxygen Species and CD4 numbers (top block) and a negative association between Oxphos/targets of Myc and CD4 numbers (middle and bottom blocks) respectively. GSEA was used to assess the association between the features and CD4 numbers/PD1+ TCM frequencies (linear regression, p<0.05). Genes that showed a significant overlap (GSEA vs inducible HIV and PD1+ TCM frequencies; significance (p-value <0.05) assessed by Fischer exact test) amongst the leading-edge gene list/pathway are represented in the heatmap (See Table S20 for overlapping gene-lists and leading-edge gene lists). Rows represent z-score normalized (red-white-blue gradient) genes and columns represent subjects from the SCOPE cohort: IR (Grey), INR-A (Red), and INR-B (Blue). The magnitude of CD4 counts (purple), frequencies of PD1+TCM (dark red) and inducible HIV (black) are plotted as annotations at the top of the heatmap. **f**, Heatmap demonstrating the association of senescent specific pathways PD1+TCM cluster. Leading-edge genes from the association of PD1+TCM and senescence pathway were summarized into a SLEA representation (z-score per pathway per sample) (See Table S20 for details). Rows represent senescent-specific pathways and columns represent samples of the SCOPE cohort: IR (Grey), INR-A (Red), and INR-B (Blue). A red-white-blue gradient is used to depict the relative SLEA score of the features, where blue represent a low relative-expression and red a high relative-expression of the feature. The magnitude of CD4 counts (purple), frequencies of PD1+TCM (dark red) and inducible HIV (black) are plotted as annotations at the top of the heatmap. **g**, Circle plot highlighting an integrated model associating leading/overlapping edges from genesets identified (classifier gene-set and biological pathways) in Figs 1 to 4, GARP+ Treg frequencies (grey), and PD1+TCM frequencies (grey) and with outcomes (CD4 counts, TILDA inducible HIV (orange)) across SCOPE cohort (n = 41). SLEA scores (z-score per pathway per sample) were calculated for each of these gene-lists (see Table S21 for the scores) and spearman’s test was used to assess correlation between all of the features mentioned. Color gradient of the Pathway node (white to red, white to blue) reflects relative enrichment (positive or negative; respectively) of pathway in Senescent-INRs compared to IR subjects. Edges between nodes represent a significant positive or negative correlation (red or blue respectively) (details for the full network are summarized in Table S21).

Lack of CD4 reconstitution was driven by senescence and impaired metabolism and as well by higher frequencies of GARP+ Tregs, increased expression of SMAD2/3 targets (including FURIN, SMOX) and heightened levels of IRF3 mediated type I IFN production and induced genes (i.e. AFF, DARC, GUK1) (Fig. 6, Supplementary Fig. 6). These pathways (IRF3 and SMAD2/3) were associated with increased frequencies of cells with inducible HIV RNA. Our integrated analysis indicates that most pathways that drive HIV persistence were negatively associated with the recovery of CD4 numbers upon cART initiation in senescent subjects. Altogether, the integration of these multi-omic results provided further evidence for the direct interplay between Treg frequencies, TGF-β production, heightened FOXO3 expression, interferon signaling, establishment of cellular senescence, impaired T cell homeostasis, and quiescent cellular and metabolic state as conditions that favor the maintenance of HIV reservoir in the unique senescent INR population described here.

## Discussion

Studies of two independent cohorts of HIV infected cART-treated subjects described in this manuscript identified cellular senescence as a mechanism that underlies HIV persistence and failure to reconstitute CD4 T-cell numbers. Our results provide a mechanistic framework where anti-inflammatory cytokines, TGF-β, VGEF and IL-13, trigger the upregulation of a transcriptional network that drives Treg differentiation. Our integrated analysis associates the increased frequencies of TGF-β producing Tregs and metabolically impaired PD-1+ TCMs with inducible HIV and poor CD4 immune reconstitution. This establishes a systemic increase in senescent cells, which is associated with HIV persistence. Notably, we demonstrate that senescence triggered by gut dysbiosis as characterized by a selective increase in the abundance of firmicutes and their metabolites including butyrates is observed only in Senescent-INRs. To our knowledge, this is the first report that provides a direct mechanistic role for the microbiome and bacterial metabolites in HIV persistence. These metabolites drive the differentiation of TGF-β producing Tregs and PD-1 expression, both of which are associated with the highest frequencies of cells with inducible HIV RNA and poor CD4 immune reconstitution.

Transcriptional profiling data underscore the critical role of signaling downstream of TGF-β that activates SMAD2/3 and the transcriptional repressor FOXO3 to induce senescence. Senescent-INRs, in our study, show increased expression of several molecules which are features of senescent cells. They include: FOXO4, a TF specifically upstream of the induction of senescence^42^; PML, triggers cellular quiescence, HIV latency and is also downstream of TGF-β; as well as molecules with anti-apoptotic activity (BCL-2, BCL-xL^119^) that ensure the survival of senescent cells. Moreover, features known to be characteristic of the senescent phenotype^68^ including decreased cell metabolism (oxidative phosphorylation), upregulation of transporters of ions/salts and inhibition of pathways that regulate cell cycle entry were characteristic of peripheral immune cell subsets in senescent INRs.

In addition to their senescent profile, these subjects are also characterized by the downregulation NF-κB and IRF7, two TFs that control pro-inflammatory pathways. The decrease in the transcriptional activity of IRF7 likely results from increased activity of the transcriptional repressor FOXO3^72^ and results in diminished expression of IRF7 target genes (including innate antivirals like CD74, TRIM4, BST2, and APOBEC3G)^71^. This impaired expression of intrinsic mechanisms of innate antiviral immunity may in part explain the increased frequencies of cells with inducible HIV in Senescent-INRs. Interestingly, while expression of FOXO3 led to poor IRF7 transcriptional activity, it was associated with an increase in IRF3 transcriptional activity. Target genes of IRF3 and FOXO3 include TFs that regulate senescence associated pathways (i.e. SOX4, a TF that inhibits T-helper cell development and is triggered by TGF-β^120^) and RIOK3, an inhibitor of NF-κB^58^. Hence, while IRF7 inhibits the anti-inflammatory activity of FOXO3^121^, IRF3 will inhibit TNF induced NF-κB activation^122^ and as well trigger the inflammasome^123^; both(decreased IRF-7 and NFκ-b) could lead to poor immune responses and heightened HIV persistence. A senescent phenotype downstream of IRF3 driven type I IFN production and signaling to induce IFN stimulated genes (ISGs) has been described in aging cells^124,125^, and could contribute to CD4 T cell senescence shown in our study. A role for type I IFN signaling and ISGs in enhancing HIV persistence was confirmed by recent studies in humanized mice whereby the blockade of IFN-α/β receptor (IFNAR) led to a reduction of HIV-1 reservoir size and to the restoration of antiviral immunity^126^. These findings are supported by the significant correlation between a type I IFN signature and HIV reservoir observed in our data. Together, this previously undescribed transcriptional cooperation between FOXO3 and IRF3 suggests novel mechanisms to understand the transcriptional regulation of senescent immune cells associated with the lack of CD4 T-cell immune reconstitution and HIV persistence.

FOXO3 is also known to transcriptionally regulate genes involved in redox balance and anti-oxidant defenses^127,128^. Our data highlights FOXO3 target gene SNCA (α-syneculin^129^), mitophagy related genes (PINK1 and PARKIN^130^) and GABARAPL2 (the autophagy regulatory gene^131^) and SLCs that are upregulated in Senescent-INRs as part of a mitochondrial damage response. The latter may help sustain processes responsible for oxidized lipid biogenesis and ROS production. Furthermore, the downregulated oxidative phosphorylation and glycolysis observed in these subjects could be driven by the upregulation of FOXO3 target genes CYB5R3 and peroxiredoxins (PRDX2, PRDX3, PRDX5) involved in maintaining redox balance and anti-oxidant defense^132^. PRDXs, specifically PRDX2, can get oxidized by ROS due to a lack of electron transport in the mitochondria^132^. Oxidized PRDX2 can then activate kinases including p38 that initiate stress responses and protect cells from ROS mediated cell death, thereby supporting the survival of senescent cells^132^. In our data, the expression of genes associated with ROS production (Catalase and superoxide dismutase) and oxidized lipid biogenesis (CD36) were positively correlated with HIV reservoir size confirming the link between senescence and HIV reservoir establishment.

The aforementioned senescence associated impaired mitochondrial metabolism was found to be enhanced in a subset of PD-1+ TCMs that were enriched in the senescent-INRs. The heightened expression of PD1+ and poor metabolic profile is suggestive of an accumulation of these cells in an early memory differentiated state specifically within Senescent-INRs. We and others have shown that PD-1 expression leads to the accumulation of cells in the early memory differentiated stage^133^ while the genetic ablation of PD-1 resulted in their increased differentiation to the effector memory stage^134^. Additionally, T-cells from Senescent-INRs show downregulation of pro-apoptotic pathways and increased expression of anti-apoptotic molecules that potentially restrain cell turnover^135^. The blockade of T-cell differentiation from TCM to TEM as a consequence of PD-1 expression could explain the low CD4 T cell numbers observed in these subjects (in contrast to TEM cells which are known to expand and proliferate thereby increasing CD4 cell numbers^112^). This limited T-cell turnover in Senescent-INRs could explain the high frequencies of cells with the multiply spiced HIV RNA, as these cells would accumulate with time.

Finally, we provide direct evidence for TGF-β producing Tregs as drivers of impaired T-cell homeostasis and HIV persistence in Senescent-INRs. In our study, the observed increase of active plasma TGF-β in Senescent-INRs is supported by the increased expression of GARP by these Tregs and activation of signal transduction cascades that lead to upregulation of FOXO3, PML and CCNT2 – all transcriptional targets of SMAD2/SMAD3 and downstream of TGF-β^35^. PML, an interferon induced anti-viral protein^136^ is triggered by TGF-β^46^ and establishes cellular quiescence by stabilizing FOXO3 and FOXO4^137^. This stabilized FOXO4 binds to p53 and plays a crucial role in the downregulation of pro-apoptotic machinery^42,43^, a major hallmark of senescent cells. *In vitro* validation experiments, in addition to molecular and cellular mechanisms described here implicate TGF-β in driving senescence and dysregulated type I IFN-driven pathways (via FOXO3 and IRF3) which impact T-cell homeostasis and promote HIV persistence. Further role of TGF-β in HIV persistence is provided by the presence of CCNT2 in the transcriptional network that is specific to senescent-INRs. CCNT2 and AFF1 are both components of the pTEFb complex which is critical for HIV replication^77,138^. Heightened levels of pTEFb in CD4 T cells could provide a mechanism to explain the higher levels of inducible HIV in these subjects.

In addition, to the direct impact of α-ketobutyrate on FOXO3 upregulation, we describe a Treg dependent induction of senescence in Senescent-INRs. Using both *ex vivo* and *in vitro* experiments, we demonstrate that the differentiation of naive CD4 T-cells into GARP+ Tregs by exposure to α-ketobutyrate and the prevalence of butyrate metabolism specific genes in Tregs. These novel findings confirm previous *in vivo* studies that have shown increased Treg differentiation upon exposure to butyric acid^106,107^. In our study, the observed increase of active plasma TGF-β in Senescent-INRs is supported by the increased expression of GARP by these Tregs and activation of signal transduction cascades that lead to upregulation of FOXO3 and PML. The elevated levels of butyrates can be attributed to an increase in ketone bodies (produced by the liver during periods of poor dietary intake^139^). These observations are supported by increased detection of both primary and secondary liver/bile acids in plasma from subjects with higher inducible HIV (i.e. senescent INRs). Alternately, the increase in plasma butyrates could also be attributed to increased bacterial metabolism^107^ and is supported by the positive association of butyrates/propionates with plasma LPS and IFABP in Senescent-INRs – providing strong support for microbial translocation as a source of these metabolites. Heightened sCD14 and increased Firmicute phylum abundance (known producers of butyrates^140,141^) in the Senescent-INRs confirms a direct interplay between microbiome/metabolic changes and establishment of the Senescent phenotype. Together these data highlight the role of butyrate induced Treg differentiation^106^ in TGF-β production which leads to subsequent CD4 T-cell senescence, lack of CD4 reconstitution and higher inducible HIV production. They reveal for the first time a mechanism downstream of commensal bacteria (exemplified by the association of the Lactobacillaceae family and alpha-ketobutyrate with inducible HIV) that can trigger the establishment of the HIV reservoir.

Our findings provide a strong rationale for the evaluation of senolytic drugs that would target senescent cells^142,143^ as promising therapeutic interventions to reduce the HIV reservoir size. Combination interventions targeting PD-1 and TGF-β, or Senolytics, may have an improved therapeutic impact in Senescent-INRs where frequencies of PD-1+ cells are a correlate of high HIV reservoir and poor CD4 T-cell reconstitution. Interestingly, senescence and TGF-β signaling have been highlighted as hallmarks of cancer where anti-PD-1 therapy has shown efficacy^144^. Our gene classifier should help identify subjects who will not respond to checkpoint therapies in HIV and cancer^145^, and could benefit from interventions that target both PD-1 and TGF-β. Senescent-INRs could also benefit from interventions that promote T-cell differentiation (e.g. IL-15) or rescue the activity of deficient metabolic pathways allowing T-cells to overcome the senescent state and help restore both CD4 T-cell numbers and cell-mediated immunity, which could decrease reservoir size^146^. The multi-omics senescent features and the classifier gene-set described here thus represent critical tools to identify Senescent-INRs. Additionally, the integrated approach used herein holds potential for the identification of novel targets and the design of optimal trials that combine different drugs/biologics targeting senescence and exhaustion to restore immune homeostasis and eradicate HIV.

### Experimental Procedures

#### Study Participants

These studies were approved by the Institutional Review Boards at University Hospitals/Case Medical Center and the Cleveland Clinic Foundation, the Vaccine and Gene Therapy Institute of Florida and University of California San Francisco; all patients provided written informed consent in accordance with the Declaration of Helsinki. ***Cohort 1 (CLIF):*** A total of 45 immunologic non-responders (INRs) and 17 immune responders (IRs) were identified from the Cleveland Immune Failure cohort. The Cleveland Immune Failure study examined immunologic indices in healthy controls and 2 groups of patients who had been receiving ART for at least 2 years with plasma HIV RNA levels below detectable levels using routine clinical assays; typically, less than 50 copies per milliliter. Transient blips in HIV RNA levels did not exclude participation if flanked by levels below limits of detection. Immune failure patients (INRs) had CD4 T cells <350/μL and immune success patients (IRs) had CD4 T cells >500/μL. Detailed clinical indices, anti-retroviral regimens and basic demographics are listed in Table S1. ***Cohort 2 (SCOPE):*** A total of 21 INRs and 20 IRs were identified from the University of California San Francisco (UCSF) SCOPE cohort. Viral suppression was defined by at least two longitudinal tests that showed plasma HIV RNA levels below the limit of detection using standard assays (<40 Abbot RealTime HIV-1 assay, <40 Roche COBAS® Ampliprep/COBAS® Taqman® HIV-1 Test, <50 branched DNA); these tests were done 3 months prior to and on the date of specimen collection. Subjects with viral blips below 200 copies/ml in the time period since beginning ART (or since the end of last treatment gap) 2 years prior to the specimen date were not excluded from the study. Basic demographics and clinical readouts at the time of collection are listed in Table S2.

#### Microarray Pre-processing and differential gene expression analyses

Whole blood was collected and lysed in RLT for RNA extraction (Qiagen, Valencia, CA, USA). For the CLIF cohort, cDNA obtained after reverse transcription reaction was hybridized to the Illumina Human HT-12 version 4 Expression BeadChip according to the manufacturer’s instruction and quantified using the Illumina iScan System. The data were extracted using the GenomeStudio software. Similarly, the SCOPE cohort samples were run using an Affymetrix microarray system. Detailed analysis of the genome array output data was conducted using the R statistical language^147^ and the Bioconductor suite^148^. Arrays displaying unusually low median expression intensity and variability across all probes relative to all arrays were discarded from the analysis. Probes that do not map to annotated RefSeq genes and control probes were removed. Quantile normalization followed by a log2 transformation using the Bioconductor package LIMMA was applied to raw microarrays intensities. To determine differences between groups or against a continuous variable, the LIMMA package^149^ was used to fit a linear model to each probe. A (moderated) Student’s *t-*test to assess significance in difference between groups, whereas a Pearson correlation test was used to assess significance against a continuous variable. The proportions of false positives were defined using the Benjamini and Hochberg method^150^. Unless indicated otherwise, genes that satisfied FDR <0.05 were selected for data mining and functional analyses.

#### Class identification, classifier training and validation

<u>*Class identification*</u> – In the CLIF cohort, 62 samples were hybridized by the Vaccine Genome Research Institute Genomics Core and 61 samples made it to the downstream analysis (44 INRs and 17 IRs) after outlier identification and removal. Initial exploratory analysis (MDS plot – Fig. 1a) followed by unsupervised analysis (heatmap of top 200 most variable probes – Supplementary Fig. 1c) revealed the discovery of two INR classes in the dataset. These presence of these subject cluster was confirmed when the Gap statistic technique^91^ was used on whole transcriptomic data to determine the optimal number of INR classes. The two classes of INRs were labeled: INR-A (an extreme INR phenotype, n = 15) and INR-B (INR phenotype proximal to IRs, n = 29). Differential expression analyses (described above) were used to identify the differences between IR and INR-A, IR and INR-B and INR-A and INR-B. *<u>Classifier training</u>* – Construction of the INR-A – INR-B Classifier (Fig. 1f-g, Supplementary Fig. 3) was done using the Prediction Analysis of Microarrays for R (PAM) package from R. PAM uses the nearest shrunken centroid methodology^151^. A standardized centroid for each class (INR-A and INR-B) was computed. Ten-fold cross-validation was used to optimize the shrunken criteria. The-shrunken criteria that resulted in the lowest crossvalidated misclassification error rate was used to generate the optimized classifier on the CLIF cohort (training cohort). Supplementary Fig. 3a represents training of the 352 features used to build the classifier on the CLIF dataset (see features in Table S11). *<u>Classifier validation</u>* – Testing the classifier on the validation set (SCOPE cohort) is illustrated in Fig. 1f-g. There are no “true” labels that define INR-A or INR-B in the SCOPE testing cohort as they both fall under the label immune non-responders. In order to assess the performance of the classifier on the validation set, the two INR groups had to be defined. An unbiased unsupervised approach similar to the one used to discover the classes in the CLIF cohort was applied on the normalized Affymetrix microarray data of the SCOPE cohort. Supplementary Fig. 3b shows a heatmap representation of the expression of the top varying probesets (top 200 variant transcripts) on the full SCOPE dataset. Two clusters of INRs were identified using unsupervised clustering (color of branches in Supplementary Fig. 3b denote the discovered classes). This discovery allowed us to label the two subgroups of INRs in the SCOPE cohort. The 352-gene classifier was then applied on the SCOPE cohort. ROC analysis was then used to assess the accuracy (80%) of the classifier on the SCOPE cohort (Fig. 1f). Throughout the study the INR classes in the SCOPE cohort were defined based on the 352-gene classifier (PCA representation of classifier genes in Fig. 1g). Whole transcriptome profiling of classifier defined classes in the SCOPE cohort is shown in Supplementary Fig. 3c.

#### Pathway analyses

*<u>Gene Set Enrichment Analysis (GSEA)</u>*^27^ throughout the study was performed to assess enrichment in pathways that discriminated between classes (Fig. 1b-d), were associated with inducible HIV (Fig. 2d and Fig. 3e), were associated with Treg cluster 7 (Fig. 3e), were associated with low CD4 counts (Fig. 4e) and were associated with PD1^hi^ROS^hi^ CD4 TCMs (Fig. 4e, f). Briefly, the whole transcriptome probe-set was collapsed to genes by assessing the most variable probes, pre-ranking of this collapsed transcriptome was done (t-statistic) and enrichment of various genesets was tested after running 1000 permutations of enrichment. *<u>Hallmark MSigDB database</u>*^25^ (v 7.0) was used to identify pathways that differentiated IR, INR-A and INR-B subjects in the CLIF cohort (Fig. 1b). This database was also used to assess pathways driving lower CD4 counts in the Senescent INRs (Fig. 4e, Table S20), frequencies of PD^hi^ROS^hi^ CD4 TCMs in the Senescent INRs (Fig. 4e, Table S20) and inducible HIV (Table S12) in the SCOPE cohort. *<u>Cell subset deconvolution</u>*: Major peripheral immune subset-specific gene expression signatures were obtained from Nakaya et. al^31^, and their enrichment was assessed (Fig. 1c, Table S7). In addition, data from a comprehensive study with sorted hematopoietic stem cell-subsets^32^ was used to generate gene-signatures specific to 38 unique subsets (each subset-specific signature was defined as genes found to be >2-fold higher with p<0.05 than in the pool of all other subsets) (Table S8). These signatures were used to assess variations in differentiated CD4 and CD8 T cell subsets in whole blood in Fig. 1d. *<u>Custom genesets</u>*: Senescence based gene signatures were extracted from Reactome database in MsigDB v7.0’s c2 module (Table S20 lists of all pathways, Fig. 4f). Genesets specific to host antiviral restriction factors (Fig. 2d, Table S12) and IFN signaling (Fig. 1b, Fig. 2d, Table S12) were extracted from Abdel-Mohsen et. al^73^ and Interferome database^76^, respectively. SMAD2/3 and HDAC1/2 genesets were also extracted from MsigDB’s c2 module by searching this module for ‘SMAD2’, ‘SMAD3’, ‘HDAC1’ and ‘HDAC2’. The results of these analyses are shown in Fig. 3e (Table S17). The enrichment of the 352-geneset classifier gene-set was done where specified (Fig. 1i, Fig. 3e, Table S17). *<u>Leading Edge overlap and Sample Level Enrichment Analyses (SLEA)</u>:* Leading edge genes from the analyses above were overlapped (significance of overlap was determined using Fischer’s Exact test; p < 0.05) to define new gene-lists that were associated with both “inducible HIV and Treg cluster 17” (Fig. 3e, Table S17) OR “CD4 counts and PD1^hi^ROS^hi^ CD4 TCMs” (Fig. 4e, Table S20). SLEA was used to generate a z-score normalized value for genelists in Fig. 2d, 3e, 4e and 4f. These scores were correlated (spearman correlation test) with each other, minor (Treg frequencies and PD1^hi^ROS^hi^ CD4 TCM frequencies) and major (inducible HIV and CD4 counts) outcomes to generate the integrated correlation network model in Fig. 5 and Table S21.

#### Over-representation, gene-mania/correlation networks and transcription Factor (TF) analyses

*<u>Over-representation</u>* of Reactome pathways in genes that are upregulated in the INR-As (vs both IRs and INR-Bs; FDR < 0.05) was assessed using the ClueGO plug-in from Cytoscape (http://apps.cytoscape.org/apps/cluego^37^), and the results of these analyses were corrected for multiple comparisons using Bonferroni step-down correction (Fig. 1e, see Table S10 for full results). *<u>GeneMania and correlations network</u>*: GeneMania Networks^152^ (http://genemania.org) were plotted to represent coexpression of genes (Supplementary Fig. 2b and Supplementary Fig. 2d). Overlap between the genes included in the networks and Gene Ontology (GO) biological process was assessed using a Fisher exact test. Correlation networks in Fig. 3d, Fig. 4d, Fig. 5a and Supplementary Fig. 5a are based on significant Spearman correlations (p-value < 0.05) between the features indicated in each figure. All networks were plotted using the Cytoscape Plug-in – specific value/category for all nodes and edges are available upon request. *<u>Transcription factor list</u>* was generated by combining genes with known ChIP-seq validated targets from CHEA^74^ (https://amp.pharm.mssm.edu/Harmonizome/resource/ChIP-X+Enrichment+Analysis) and ENCODE (https://amp.pharm.mssm.edu/Harmonizome/dataset/ENCODE+Transcription+F_actor+Targets) databases. The expression of these TFs was tested in INR-As vs INR-Bs or IRs (Supplementary Fig. 2c). IRF3 targets and FOXO3 targets were extracted from the ENCODE and CHEA databases cited above, respectively. Coexpressed targets of FOXO3 and IRF3 that correlated with inducible HIV are represented in Fig. 2d (Table S12).

#### Cell Preparation and flow cytometry

PBMCs were prepared from whole blood by ficoll-hypaque density sedimentation and cryopreserved in 10% dimethyl sulfoxide and 90% FBS. Two panels to evaluate Treg function and mitochondrial dysfunction in memory CD4s were run using previously titrated antibodies summarized in Table S13, S14 and S18. The cells were surface stained for 20 minutes in the dark at room temperature, washed, fixed and permeabilized using the eBioscience™ Foxp3 / Transcription Factor Staining Buffer Set (Cat# 00-5523-00), as per manufacture instructions. Intracellular staining was performed in Perm-Wash provided by the kit for 45 minutes at 4°C. Samples were washed and re-suspended in staining buffer for acquisition. ~500,000 live-gated events were collected per sample on the LSRII flow cytometer (Becton Dickinson, San Jose, CA). Initial data cleaning and pre-gating was done using the Flow-Jo X software (TreeStar, Ashland, OR) (Supplementary Fig. 6). Briefly, lymphocyte gate based on FSC-A and SSC-A was defined. Single cells were then selected using a FSC-A x FCS-H gate. Live CD3+CD4+CD8-cells were gated and exported for unbiased clustering analyses (Supplementary Fig. 6). For both panels, projection of the density of cells expressing markers of interest (Table S14, Table S18) were visualized/plotted on a 2-dimensional UMAP (https://arxiv.org/abs/1802.03426, https://github.com/lmcinnes/umap). Clusters of cells using the RPhenograph package after (https://github.com/jacoblevine/PhenoGraph) after concatenating all samples per panel and bi-exponentially transforming each marker (Fig. 3a and c, Fig. 4a). Differences in cluster frequencies per group and MFI for each marker per cluster, for each are shown in Table S14 and Table S18.

#### Measurement of inducible reservoir

The HIV reservoir was measured using the tat/rev induced limiting dilution assay (TILDA). This method measures the inducible reservoir by RT-qPCR using the limiting dilution assay previously described^70^. Briefly, 2 x10^6^ CD4 T cells from 20×10^6^ PBMCs were enriched using a CD4 T cell negative selection kit (Stem Cell). These cells were stimulated with PMA/Iono (100ng/ml and 1μg/ml, respectively) for 12 hours. A total of 744,000 stimulated cells were then distributed in a limiting dilution in a 96-well plate and tat/rev msRNA expression was directly quantified (without prior RNA extraction) by semi-nested real-time PCR^70^. The frequencies of cells positive for inducible tat/rev msRNA per 10^6^ CD4+ T cells were determined by using the maximum likelihood method (http://bioinf.wehi.edu.au/software/elda) (Fig. 1h,i).

#### Measurement of plasma biomarkers

Plasma IL-6 in the CLIF cohort was measured using a high sensitivity ELISA kit for human IL-6 (Quantikine HS) from R&D Systems (Minneapolis, MN). Plasma levels of IL-7 in the CLIF cohort were measured by high sensitivity IL-7 ELISA (Quantikine HS, R&D Systems, Minneapolis, MN). Plasma levels of IP10 in the CLIF cohort were measured by IP-10 ELISA (Quantikine, R&D Systems, Minneapolis, MN). Plasma levels of human Intestinal Fatty Acid Binding Protein (I-FABP) in the CLIF cohort were measured using a DuoSet ELISA Development kit from R&D Systems (Minneapolis, MN) following the manufacturer’s protocol. Levels of D-dimers in the CLIF cohort were measured using the Asserachrom D-DI immunoassay (Diagnostica Stago, Asnieres France) (Supplementary Fig 1. d-m). *<u>Measurement of LPS in the CLIF cohort</u>*: Plasma samples were diluted to 10% or 20% with endotoxin free water and then heated to 85°C for 15 minutes to denature plasma proteins. We then quantified plasma levels of LPS with a commercially available Limulus Amebocyte Lysate (LAL) assay (QCL-1000, Lonza, Walkersville, MD) according to the manufacturer’s protocol. <u>*Multiplex ELISA (Mesoscale):*</u> U-PLEX assay (Meso Scale MULTI-ARRAY Technology) commercially available by Meso Scale Discovery (MSD) was used for plasma cytokine detection. The assay was performed according to the manufacturer’s instructions (https://www.mesoscale.com/en/technical_resources/technical_literature/techncal_notes_search). 25μL of plasma from each donor was combined with the biotinylated antibody plus the assigned linker and the SULFO-TAG™ conjugated detection antibody; in parallel a multi-analyte calibrator standard was prepared by doing 4-fold serial dilutions. Both samples and calibrators were mixed with the Read buffer and loaded in a 10-spot U-PLEX plate, which was read by the MESO QuickPlex SQ 120. The plasma cytokines values (pg/mL) were extrapolated from the standard curve of each specific analyte. Cytokine clustering (Supplementary Fig. 4a,b,c) was performed using independent methods: (gap statistic method to identify and characterize optimal number of k-means clusters, and hierarchical clustering (ward clustering; Euclidean distance^91^).

#### Microbiome and Metabolome

*Pathogen-Sequencing*: Host microbiomes were analyzed using the PathSeq technology. PathSeq involves deep sequencing of fragmented RNA and DNA (~100-1000 bases) in plasma to obtain an overall quantitative and qualitative measure of the viral, bacterial, fungal, parasite and helminth burden. Following RNA-sequencing, the reads that map onto host genomic and transcriptomic regions were identified and removed from the dataset. De novo assembly is then performed on the remaining, non-host sequences. The assembled contigs were aligned against the comprehensive “nt” database from NCBI to determine the microorganisms for which assembled sequences were derived. We used the Bray-Curtis dissimilarity statistic ^110^ (Supplementary Fig. 5a,b) to assess the beta diversity and assess differential distribution of the abundance or total read counts of all phyla to species in the microbiome PathSeq data. *Metabolomics:* In collaboration with Metabolon, plasma metabolite levels for up to 1300 metabolites were measured. Metabolon uses four ultra-high-performance liquid chromatography/tandem accurate mass spectrometry (UHPLC/MS/MS) methods in a highly controlled environment to reduce noise and produce accurate results. The data generated using UHPLC/MS/MS were referenced against a well-established library of known and novel metabolites.

#### Treg differentiation in the presence of alpha-ketobutyrate

Naïve CD4 T cells extracted from 8 healthy donors were isolated using the EasySep™ Human Naïve CD4+ T Cell Isolation Kit (StemCell Technologies Catalog# 19555) and differentiated in vitro using the protocols described by Rudensky^106^. Briefly, 100-200,000 naïve CD4 T cells were stimulated with Dynabeads Human T-Activator CD3/CD28 beads (ThermoFisher Scientific Cat# 111.31D, 1 bead/3 cells), 100 U/mL of IL-2 (R&D Systems Cat# 202-IL-500), 0 to 1ng/mL of TGF-β (R&D Systems Cat# 240-B-010) and alpha-ketobutyrate (Cat#, from 0mM to 5mM) for 3 days in 96-well round bottom plates. Viability post-stimulation, assessed poststimulation by staining with a fixable viability dye, was found to be >80% in all stimulation conditions. Secreted cytokines in the supernatants were quantified using the Mesoscale Discovery platform/kits described above. CD4 T cells were intracellularly stained with the Treg phenotyping panel described (Table. S8).

#### Latency establishment and reactivation assay

The latency and reactivation assays developed by our group^112^ was used to assess the latency establishment in memory CD4 T cells post-stimulation with increasing doses of TGF-β. Briefly, memory CD4 T cells spinoculated with HIV (HIV strain information listed^112^) were incubated in the presence of antiretrovirals (effavirenz, Saquinavir and Ralegravir) and latency establishment media (supplemented with 0-40ng/mL of TGF-β) for 13 days (or when the frequencies of HIV-p24+, measured by flow cytometry, CD4 T cells was negligible). Integrated HIV DNA was measured at this stage using the protocol described^112^. Reactivation of HIV post-latency was done after stimulation with 1ug/mL anti-CD3 and 1ug/mL of anti-CD28 for ~48 hours; reactivated cells were quantified by monitoring the frequencies of p24+ cells using flow cytometry.

#### Other statistical analyses

All univariate group-differences were analyzed using a non-parametric Wilcoxon-ranked test. All univariate correlation analyses were done using a non-parametric Spearman’s test. P<0.05 is reported as significant.

#### Full Data Availability Statement

The raw and normalized data matrices for gene expression analysis of the CLIF and SCOPE cohorts were deposited into the Gene Expression Omnibus (GEO) database; GSE143742.

#### Financial support

This work was supported by the National Institutes of Health (grants UO1 AI 105937 and RO1 AI 110334 and RO1 AI 11444201), the CWRU Center for AIDS Research (grant AI 36219), DARE (U19 AI 096109), and the Fasenmyer Foundation. Rafick-Pierre Sekaly is the Richard J. Fasenmyer Professor of Immunopathogenesis.

## Supplementary Figures (S1-S7)

**Supplementary Figure 1.**
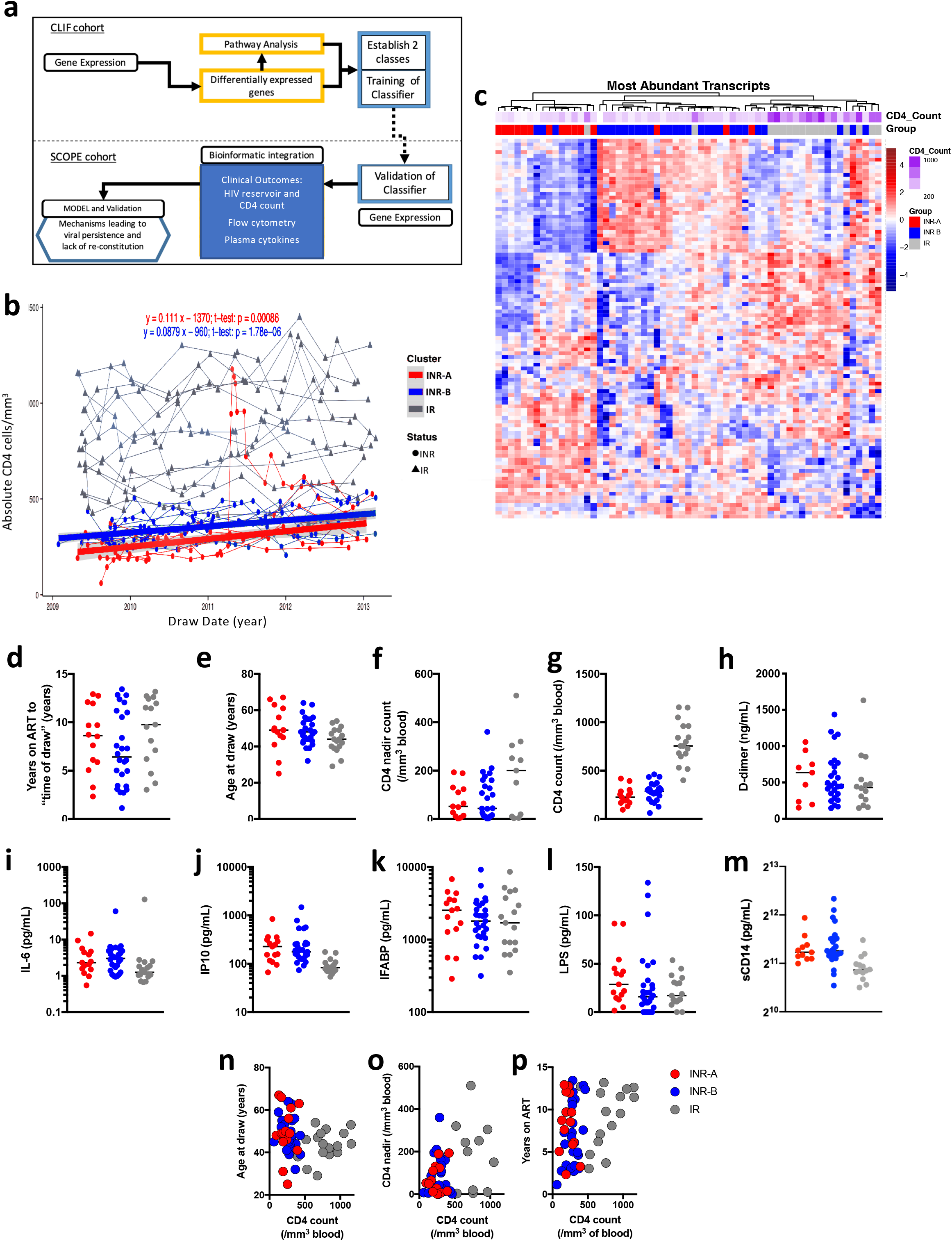
Gene Expression profiling provides a unique tool to identify two heterogeneous INR subject groups. **a**, Flow chart illustrating the steps in the multi–omics data analyses and integration utilized across two independent cohorts of HIV-infected ART-treated subjects. **b**, INR groups remain relatively stable over time and show slow CD4 T cell reconstitution. Plot represents absolute CD4 counts over 3 years for all samples in each group. X-axis represents the draw date (in years) and y-axis represents the absolute CD4 count in cells/mm3. Red = INR-A, Blue =INR-B, Grey= IR. The fitted lines represent the rate of CD4 reconstitution over time (y=mx + b). **c**, Unsupervised analysis of gene expression data identifies three groups of HIV subjects treated with ART (IR, INR-A and INR-B) in the CLIF cohort. Gap Statistics revealed n=2 optimal INR clusters (bootstrap n = 500, p= 0.00293). Heatmap shows the expression of the top 200 genes that distinguish the 3 groups. Hierarchical clustering (Euclidian distance, complete linkage) was used to re-group samples with similar expression of the top 200 varying genes. Gene Expression is represented as a gene-wise standardized expression (Z-score). **d-o**, several parameters associated with poor immune reconstitution in ART-treated subjects including: Duration of ART treatment (d), Age (e), CD4 Nadir (f) and CD4 counts at draw (g), Levels of D-dimer (h), Levels of IL6 (i), IP10 (j), IFABP (k), LPS (l), sCD14 (m) were assessed for their ability to distinguish the two groups of ART-treated subjects. A Welch t-test was used to assess significance. INR-As and INR-Bs could not be distinguished for CD4 numbers and age (m), CD4 Nadir (n), or Years on ART (o). A Spearman correlation was used to assess significance.

**Supplementary Figure 2.**
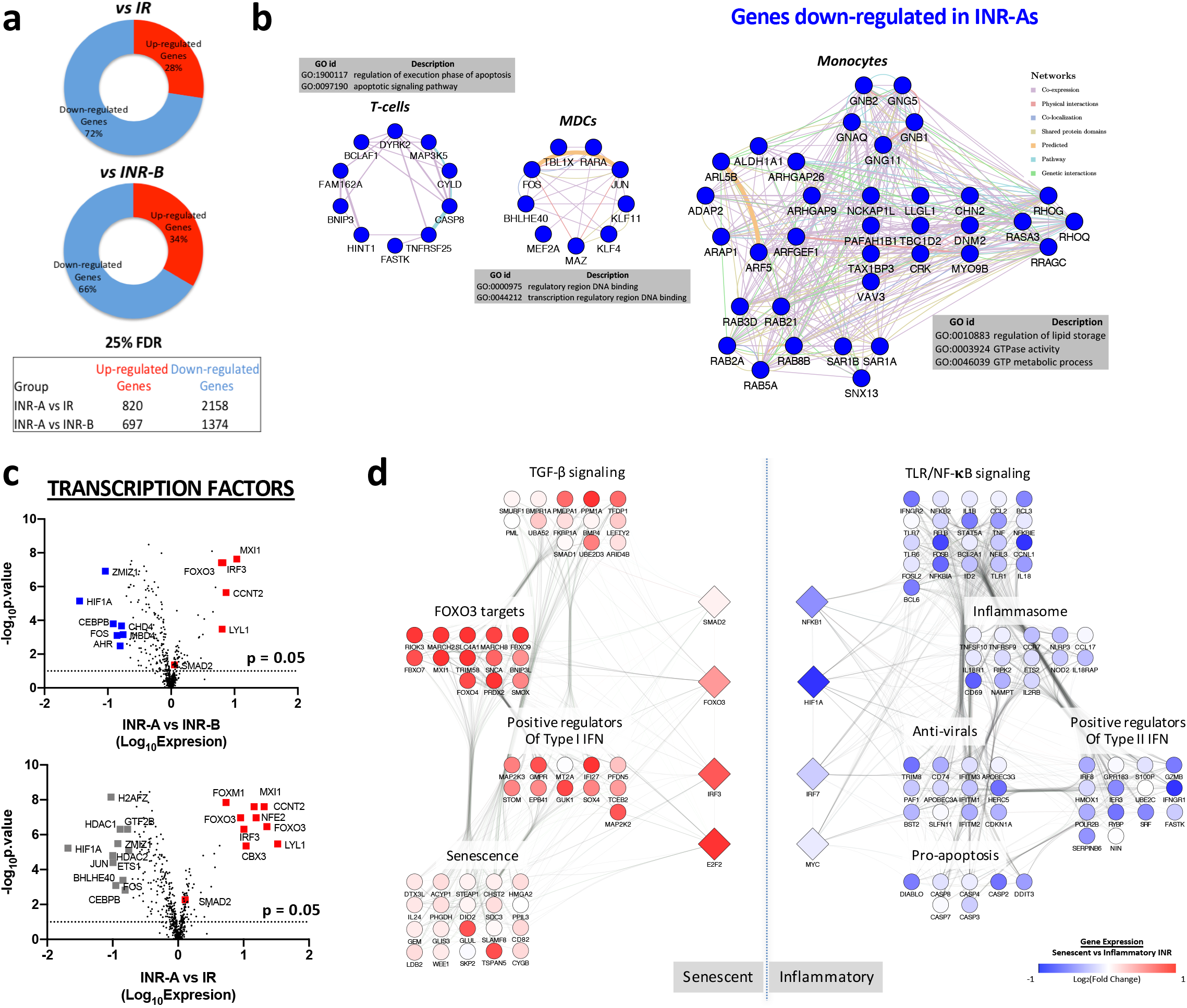
Diminished global transcriptional activity within INR-A subjects maps to down-regulation of T-cell, mDC and monocyte subset specific gene signatures and lack of effector CD8 function. **a**, Pie charts illustrating the majority of the transcriptional signal (~70%) specific to Senescent-INR subjects is due to down-regulation of genes. The majority of genes differentially expressed between INR-As and IRs (left) or INR-As and INR-Bs (right) are down-regulated (details of DEGs are in Table S3-S5). **b**, Networks highlighting the genes and functional annotations of T-cell, MDC, and Monocyte specific gene expression signatures down-regulated in INR-A subjects. GSEA (p<0.05) were used to assess significance of the down-regulation of subset genesets, Gene Ontology (GO) was used to infer the biological function (overrepresentation test p<0.05) and Gene Mania was used to plot co-expression networks. **c**, Volcano Plots illustrating the top Transcription Factors (TFs, squares) up- and down-regulated in INR-A vs IR (top panel) and INR-A vs INR-B (bottom panel) subjects. X-axis represents the log_10_FC of TF expression, and Y-axis represents the -log_10_(p-value); dotted lines highlight the p-value ≤ 0.05 cutoff. Top TFs up-regulated in INR-As are colored in red, whereas TFs specific to IRs or INR-Bs are colored in grey or blue respectively (See Table S9 for details). **d,** A functional network of the core TFs (diamonds) and their target genes (circles) up-regulated in INR-A subjects (red, left portion) or INR-Bs (blue, left portion). Network Inference and Cytoscape were used to plot the network. Reactome and GO terms were used to annotate the biological functions of the top downstream targets of the TFs up-regulated in INR-A subjects. As indicated in the figure, the node color represents log2 fold change in expression between the two INR classes. Edges within the network depict a link between a TF and its targets.

**Supplementary Figure 3.**
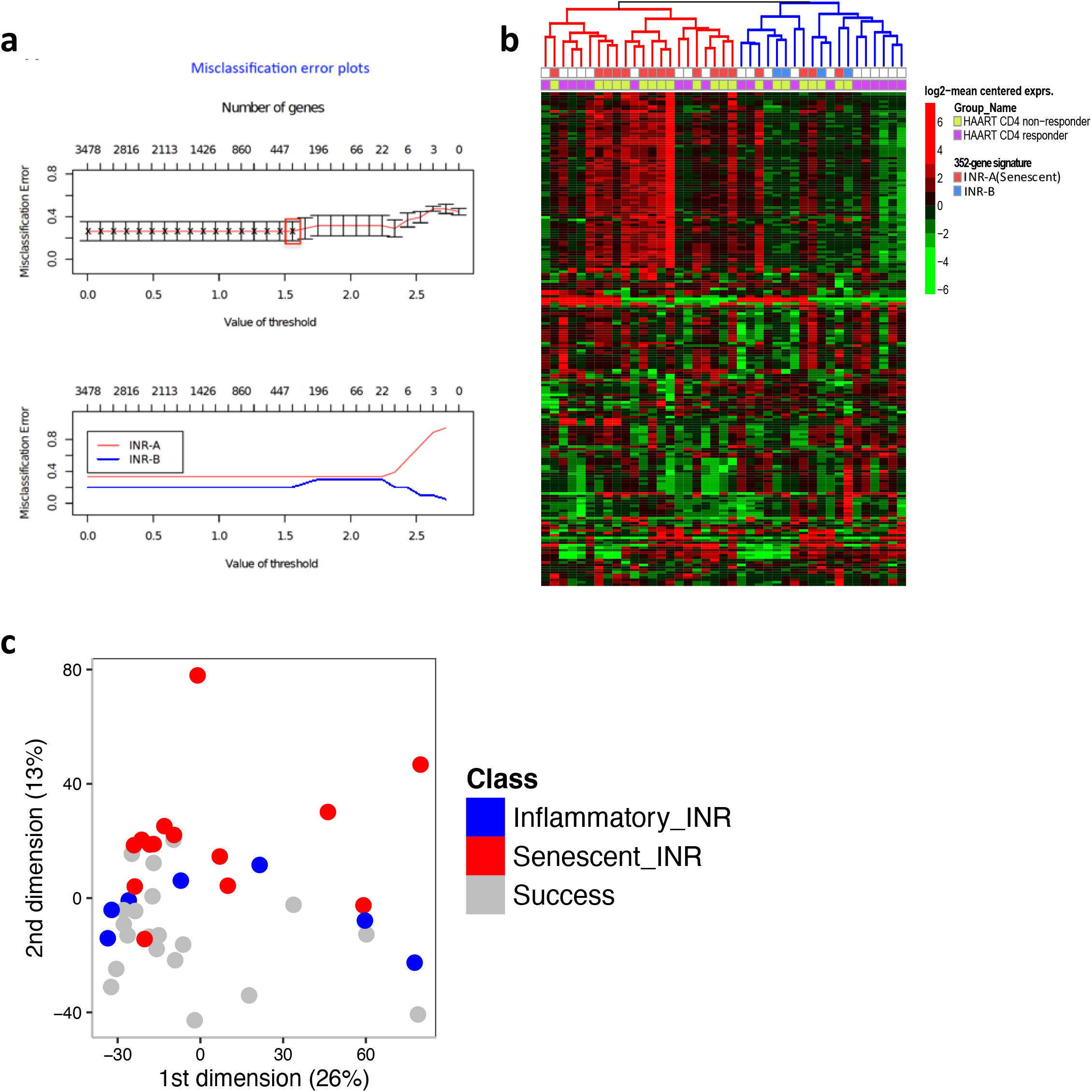
Construction and Validation of the gene-based classifier. **a**, Training of the 352-gene classifier on the CLIF cohort. Results of the 10-fold cross validation. Misclassification Error plot represents the optimal number of features (genes) that corresponds to the lowest misclassification error rate. The pamr package in R was used to train the classifier via the nearest shrunken centroid method; 352 genes were selected to segregate the two groups of immune non-responders (See Table S11 for list of classifier genes). **b**, Heatmap representation of the expression of the top 200 varying probesets in the SCOPE cohort. The expression intensities are represented using green-black-red color scale. Rows correspond to probesets and columns correspond to the profiled samples. Hierarchical clustering based on a complete linkage was used to regroup samples with similar gene-expression profiles. Two clusters of INRs (depicted in red and blue on the dendrogram) were identified using the unsupervised clustering. **c**, Multidimensional scaling (MDS) plot highlights the overall transcriptomic variation within the SCOPE validation cohort.

**Supplementary Figure 4.**
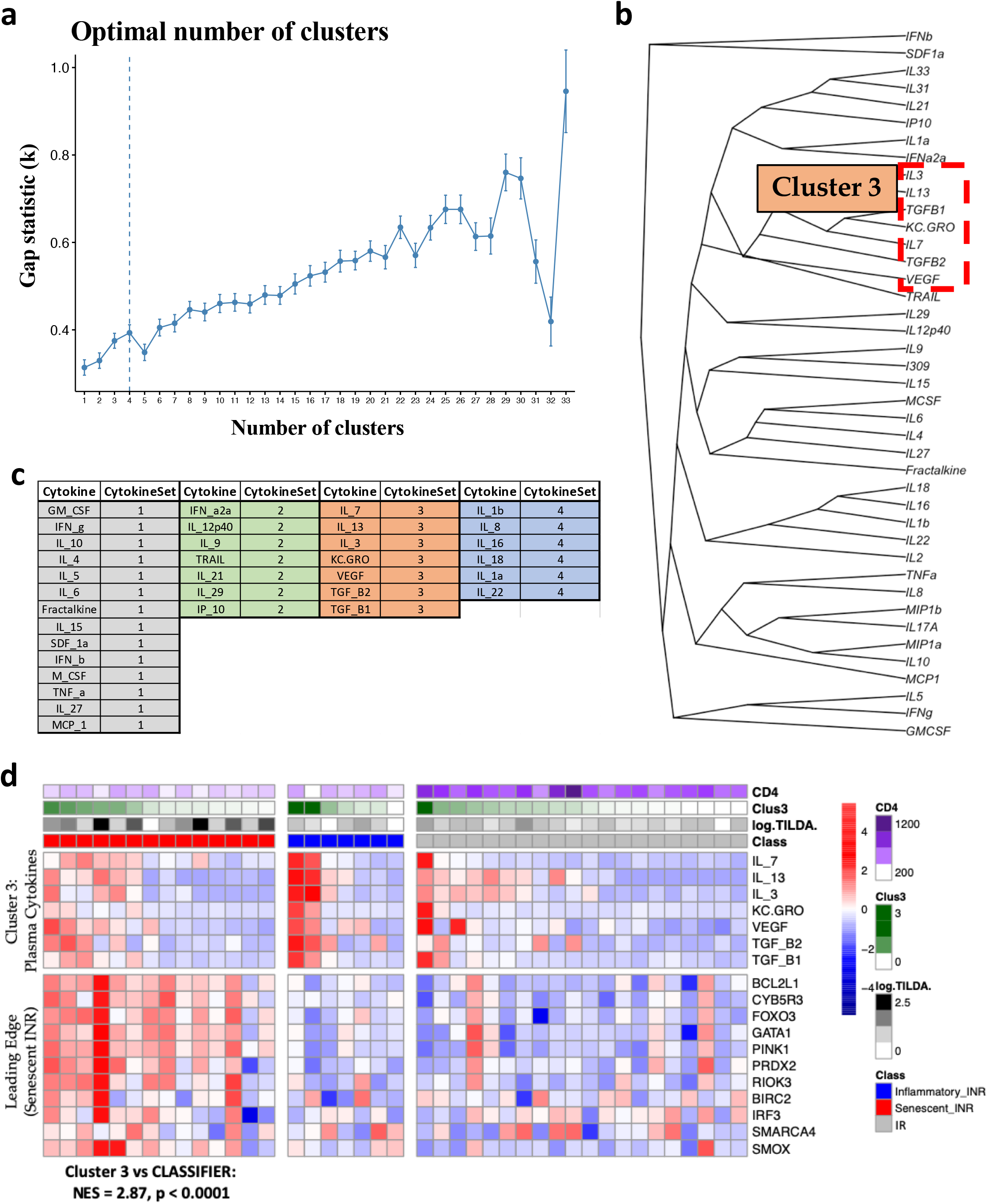
Identification of a specific Senescent Cytokine Cluster. **a**, K-means clustering and the GAP statistic technique^91^ were used to identify the optimal number of cytokine clusters after 42 cytokines were measured using the meso-scale platform. Four clusters were observed. **b**, Hierarchical clustering technique identified similar clusters. Two stable clusters (identified concurrently by k-means and hierarchal clustering) and were observed, one of which contained TGF-β1/2, IL3/7/13, KC.GRO and VEGF. **c**, Table summarizes the cytokine Cluster members identified in (a). **d**, Heatmap demonstrating the expression of the TGF-β1/2 driven cytokine cluster “Cluster 3” (top portion) and the expression of classifier features correlated to Cluster 3 (bottom portion) across all subjects of the SCOPE cohort. Rows represent the features (plasma cytokine levels or TF expression levels) and columns represent samples of the SCOPE cohort: IR (Grey), INR-A (Red), and INR-B (Blue). A red-white-blue gradient is used to depict the relative expression of the features, where blue represent a low relative-expression and red a high relative-expression of the feature. The magnitude of the Senescent Cytokine Cluster (Cluster 3, dark green) is plotted as annotations at the top of the heatmap. GSEA was used to assess the association between the classifier features and Cluster 3 of senescent cytokines (linear regression: classifier genes ~ Cluster 3, GSEA: NES=2.87, p<0.01), and the leading-edge features are represented (See Table S15 for detailed list of genes and associations of cytokine cluster 3 with transcriptome of all groups).

**Supplementary Figure 5.**
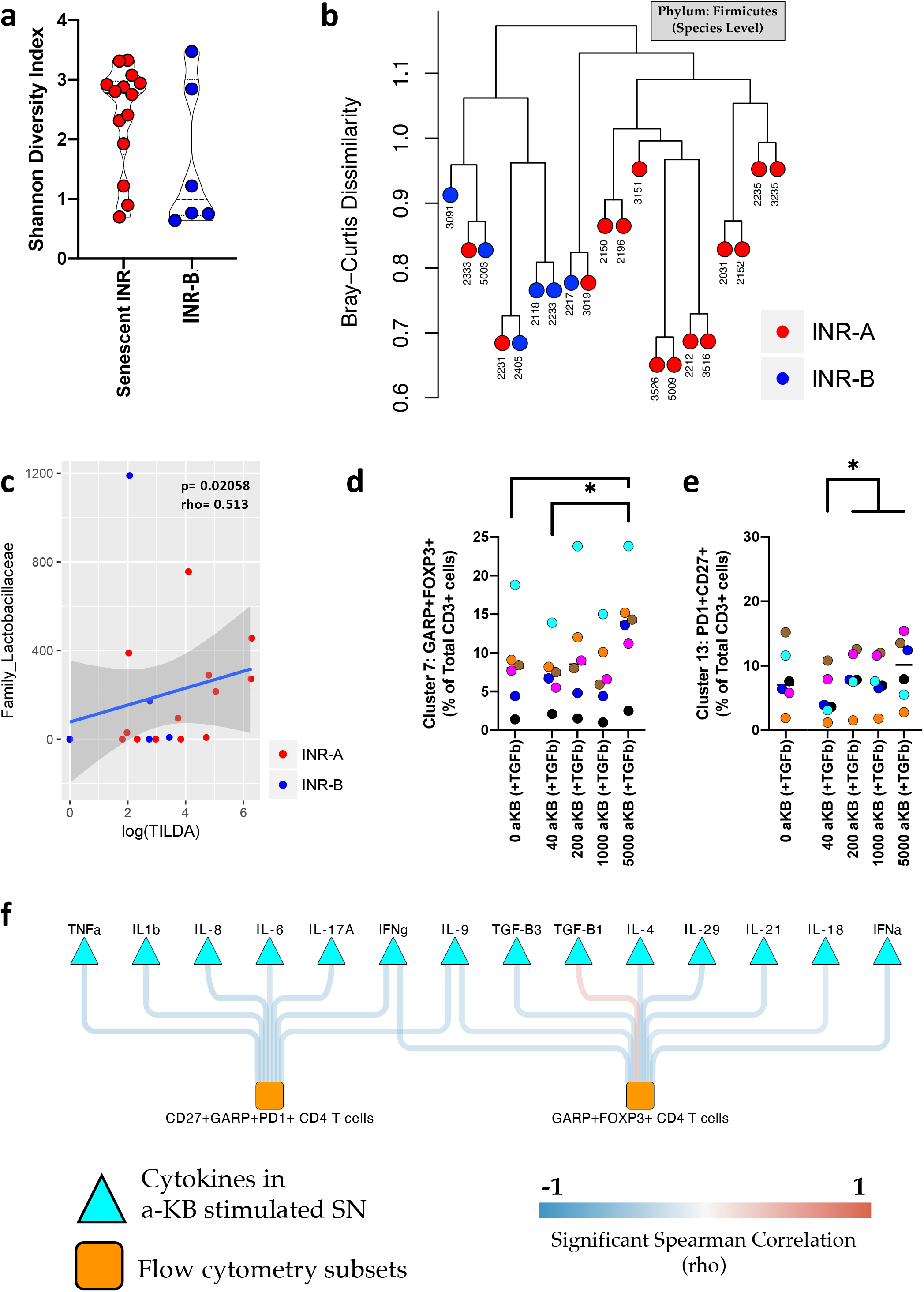
Microbiome and Metabolome differences associated with Senescent-INR subjects map to Firmicutes and Butyrates. **a**, Violin jitter plot highlighting differences in PathSeq species diversity between Senescent-INR subjects (red) and inflammatory-INR subjects (blue) of the SCOPE cohort. The Shannon diversity index (H) accounting for both abundance and evenness of the species present was used and plotted along the y-axis. **b**, Bray-Curtis dissimilarity statistic to assess the beta diversity of the Firmicute species abundance in the microbiome PathSeq data of the SCOPE cohort. Clustering dendrogram reflects differential distribution of Senescent-INR (right most branch) and INR-B (left-most branch) subjects summarizing a difference in the Firmicute microbial composition between these subjects. Circles reflect subjects: Senescent-INR (Red), and INR-B (Blue). Clustering distances reflect similarities and differences at the species level. **c**, Scatter plot highlighting a positive association between frequencies of CD4 T cells with inducible HIV as measured by TILDA plotted along the X-axis and levels of a specific family of the Firmicute phylum: Lactobacillaceae plotted along the Y-axis in -INR (red) and INR-B (blue) subjects of the SCOPE cohort. **d-e**, Jitter plots highlighting that increasing concentrations of alpha-ketobutyrate preferentially led to differentiation into GARP+FOXP3+ cells (**d**) and increased levels of PD1 expressing quiescent cells (**e**). GARP+FOXP3+ and PD1+CD27+ levels plotted along the Y-axis, while alpha-ketobutyrate increasing concentration levels are indicated on the X-axis. Conditions in the presence of TFG-b are designated with (+TGF-β). A Mann-Whitney U-test was used to assess significance across concentrations (* represents p-value <0.05 between groups). **f**, CD4 T cell subsets enriched after stimulation with alpha-ketobutyrate (i.e. GARP+ Tregs and PD1+ TCM; nodes in orange squares), were significantly positively associated (p < 0.05; Spearman’s rho>0 – red edges) with an increase in secreted TGF-β1 and a significant decrease in effector cytokines like IL17A, IFNg and IL9 (p < 0.05; Spearman’s rho<0 – blue edges).

**Supplementary Figure 6.**
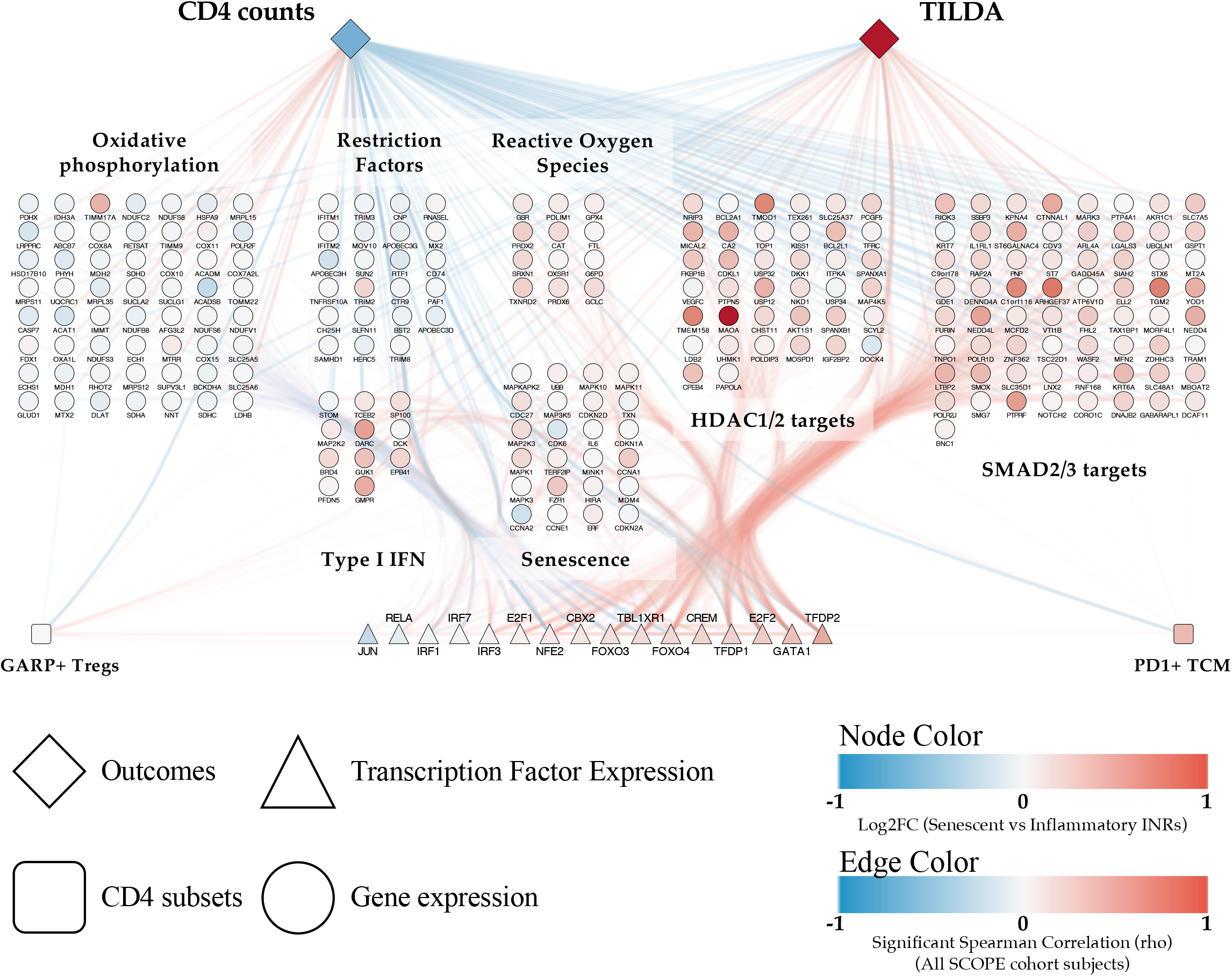
A multi-omic gene expression and flow cytometry model correlates with the magnitude of CD4 counts and inducible HIV. Detailed network depicting an integrated model highlighting the association between gene-expression outcomes (genes and pathways, circular nodes), flow cytometry readouts (square nodes) and clinical outcomes (diamond nodes). Circular nodes reflect leading edge genes of biological pathways listed and are colored by the fold change in Senescent INR subjects. Triangular nodes reflect highlight transcription factors and are colored by the fold change in Senescent INR subjects. Edges between nodes represent a significant positive or negative correlation (red or blue respectively). A spearman correlation was used to assess significance (p-value < 0.05).

**Supplementary Figure 7.**
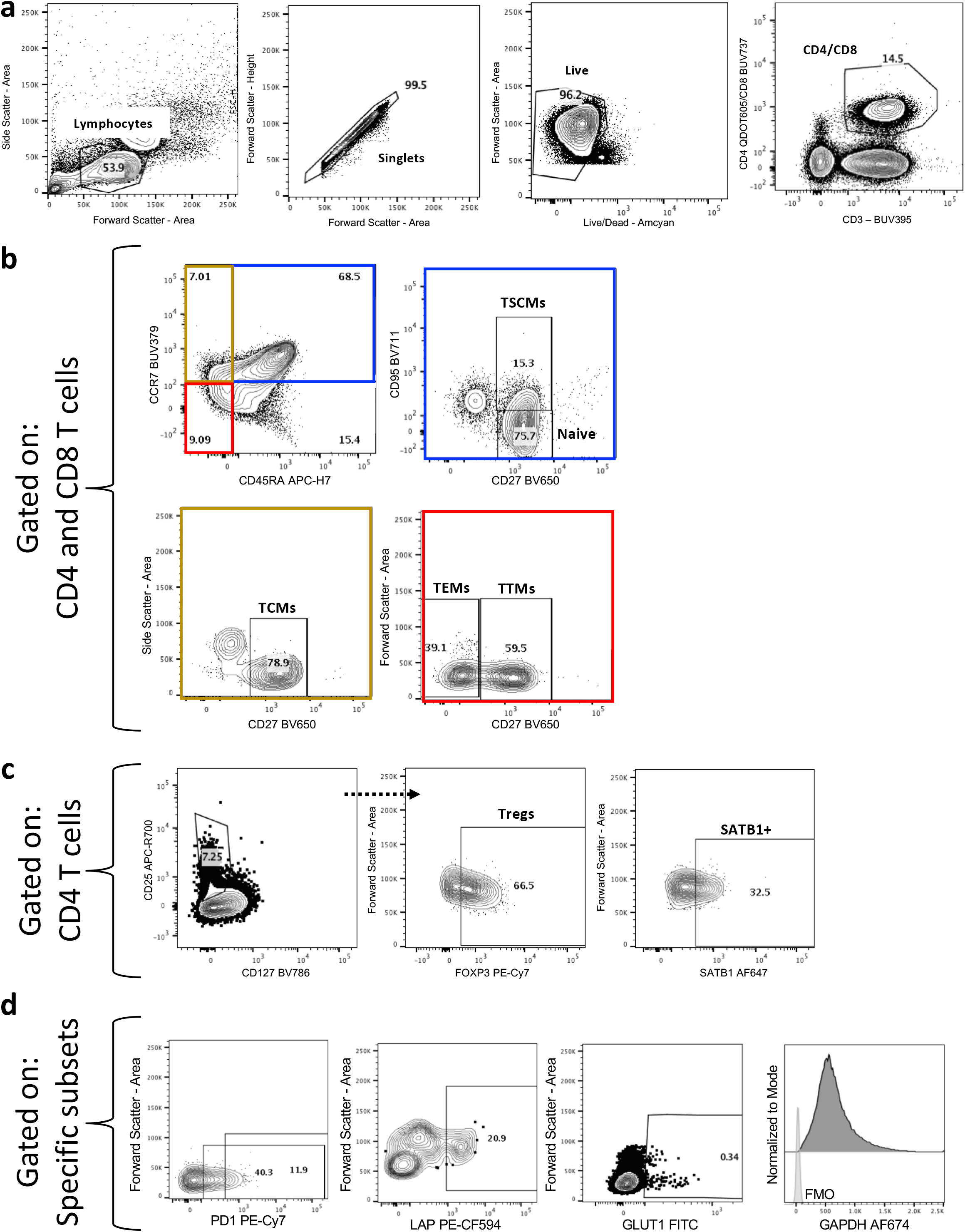
Flow cytometry gating strategy for the markers used in this study. **a**, A lymphocyte gate based on FSC-A and SSC-A was defined. Single cells were then selected by FSC-A x FCS-H gate. Live cells were gated and successive gates to define T cell populations (CD3+CD4+ and CD3+CD8+) were made. **b**, Memory phenotypes gating of CD4 and CD8 subsets (Tcm: CD45RA-CCR7+CD27+, Ttm: CD45RA-CCR7-CD27+, Tem: CD45RA-CCR7-CD27-, Naive: CD45RA+CCR7+CD27+CD95- and Tscm: CD45RA+CCR7+CD27+CD95+) were performed. **c**, Tregs were identified after initial gating on CD4 T cells and the expression of SATB1 was measured in this subset. **d**, the frequencies of PD1+, LAP+, a surrogate marker for TGF-β expression and GLUT1+ cells were measured in all subsets using the gating strategy described here.

